# ADAR2 deaminase activity promotes Th17 effector function and protects against intestine inflammation

**DOI:** 10.1101/2020.09.22.308221

**Authors:** Shengyun Ma, Yajing Hao, Benjamin S. Cho, Nicholas Chen, Anna Zheng, Shuyang Zhang, Ge Sun, Parth R. Patel, Yuxin Li, Brian A Yee, Gene W Yeo, Bing Zhou, Xiang-Dong Fu, Wendy Jia Men Huang

## Abstract

ADAR1 and ADAR2 catalyze adenosine-to-inosine (A-to-I) editing, the most common post-transcriptional modification in RNA. While ADAR1 is ubiquitously expressed and plays a critical role in preventing activation of the host immune system, ADAR2 exhibits tissue-specific and inducible expression patterns, and its function in the immune system is not known. Here, we identify an intragenic super-enhancer involved in the dramatic induction of ADAR2 when naïve helper T cells differentiate toward the Th17 lineage. By editing the inverted repeat sequences at the 3’ untranslated region (UTR) of *Malt1*, which encodes a component of the NF-κB activation complex, ADAR2 promotes MALT1 expression and Th17 effector function. Interference with the ADAR2-MALT1 pathway dampens the production of Th17 cytokines and promotes T cell-mediated colitis. This study expands our understanding of RNA editing in adaptive immunity and identifies the ADAR2-MALT1-IL-17A axis as a potential therapeutic target for inflammatory conditions in the intestine.

## INTRODUCTION

T helper 17 (Th17) lymphocytes residing in the gastrointestinal tract regulate local tissue homeostasis and response to microbial challenges (*1, 2*). These cells produce a signature cytokine called interleukin-17A (IL-17A) (*3*), which is essential for epithelial barrier function (*4–6*) and anti-bacterial and anti-fungi defense (*7–9*). Genetic polymorphisms of *Il17a* have been linked to inflammatory bowel disease (IBD) susceptibility (*10*). Neutralization of IL-17A or its receptor worsens intestine inflammation in IBD patients (*11, 12*), suggesting a protective role of the IL-17A axis in the gut.

Th17 differentiation is induced by IL-6, TGFβ, IL-21, IL-23, and T cell receptor (TCR) signaling (*13*). TCR activation promotes oligomerization of the CBM complex, containing the Caspase recruitment domain-containing protein CARMA1 (CARD11), B cell lymphoma-10 (BCL10), and Mucosa-associated lymphoid tissue lymphoma translocation protein 1 (MALT1) proteins (*14*). The CBM complex acts as a scaffold for the recruitment of effector proteins involved in NF-κB activation (*15*). MALT1-mediated cleavage of RelB (*16*), A20 (also known as TNFAIP3) (*17*), and cylindromatosis (CYLD) further modulates the NF-κB and JNK signaling cascades (*18*). Genetic and pharmacologic studies show that MALT1 is essential for Th17-mediated anti-fungal immunity and encephalomyelitis pathogenesis (*19, 20*). But the molecular pathways regulating MALT1 levels in T cells and its role in IBD remain unclear. In this study, we report a surprising finding that MALT1 expression in Th17 cells is regulated by an RNA modification-dependent mechanism.

One of the most abundant RNA modifications in the mammalian transcriptome is the deamination of adenosine to inosine (A-to-I editing), which is catalyzed by the Adenosine Deaminase Acting on RNAs (ADAR1 and ADAR2) (*21, 22*). Inosine is recognized by cellular machinery as guanosine, therefore A-to-I editing has been reported to impact gene expression at the level of pre-mRNA splicing, RNA stability, RNA localization, microRNA targeting, and/or protein translation (*23*). ADAR1 is ubiquitously expressed, and it is well appreciated for its role in editing repetitive elements to prevent aberrant activation of the dsRNA sensor melanoma differentiation-associated protein 5 (MDA5) and subsequent interferon responses and inflammation (*24–26*). In the adaptive immune system, T cell-specific deletion of *Adar1* impaired negative selection in the thymus, resulting in the development of an autoimmune condition in the intestine (*27*). Unlike ADAR1, ADAR2 shows restricted basal expression. It is highly expressed in the brain under homeostatic conditions (*27, 28*) and is best known for modulating the coding potential of the glutamate receptor subunit GluA2 essential for neuronal activities (*29*). Although ADAR1 role in innate immunity is well-documented, the role of ADAR1, ADAR2, and A-to-I editing in T cell differentiation and function are largely unknown.

Here, we report that naïve helper T cells polarizing toward the Th17 lineage undergo a switch in ADARs expression, resulting in a global shift in the A-to-I editome. This process is evolutionarily conserved between mice and humans. In murine Th17 cells, elevated ADAR2 levels are maintained by an intragenic super-enhancer. One target of ADAR2 is the MALT1 transcript essential for Th17 effector function. ADAR2 edits a set of highly selective sites on the B2 SINE repetitive elements present in the *Malt1* 3’ untranslated region (UTR) to promote protein translation. Genetic alteration of this pathway reduces Th17 cytokine production and tissue repair capacities during colitis. These findings reveal an essential role of ADAR2-mediated RNA editing in T cell and mucosal biology.

## RESULTS

### ADAR1 and ADAR2 are dynamically regulated during T helper cell differentiation

RNA binding proteins (RBPs) regulate mRNA processing, stability, transport, and translation (*30*). In CD4^+^ T helper cells, aside from the known roles of Ago1/Ago2 (*31–33*) and ZFP36 (*30*), the functions of other abundantly expressed RBPs have not been investigated. Of the RBPs curated by ATtRACT database (*34*), we detected 170 RBPs abundantly expressed in T helper cells, 72 of which displayed dynamic expression patterns upon naïve cells differentiating into the Th17 lineage (Figure 1A and Table S1). Compared to naïve cells, 29 (40%) RBP-encoding transcripts were downregulated and 43 (60%) upregulated in Th17 cells. Among those upregulated RBPs in Th17 cells, *Adarb1*, which encodes ADAR2, was the most significantly increased (Figure 1A). An independent RNA-seq dataset from a previous report also confirms these observations (*35*), showing the highest induction of *Adarb1* in Th17 cells among all CD4^+^ T cell subsets (Figure S1A). In contrast, the levels of *Adar* transcripts for ADAR1 were lower in Th17 cells compared to naïve cells (Figure 1A). The distinct expression patterns of these adenosine deaminases suggest that they serve differentiation stage-related and/or subset-specific functions in helper T cells.

**Figure 1.**
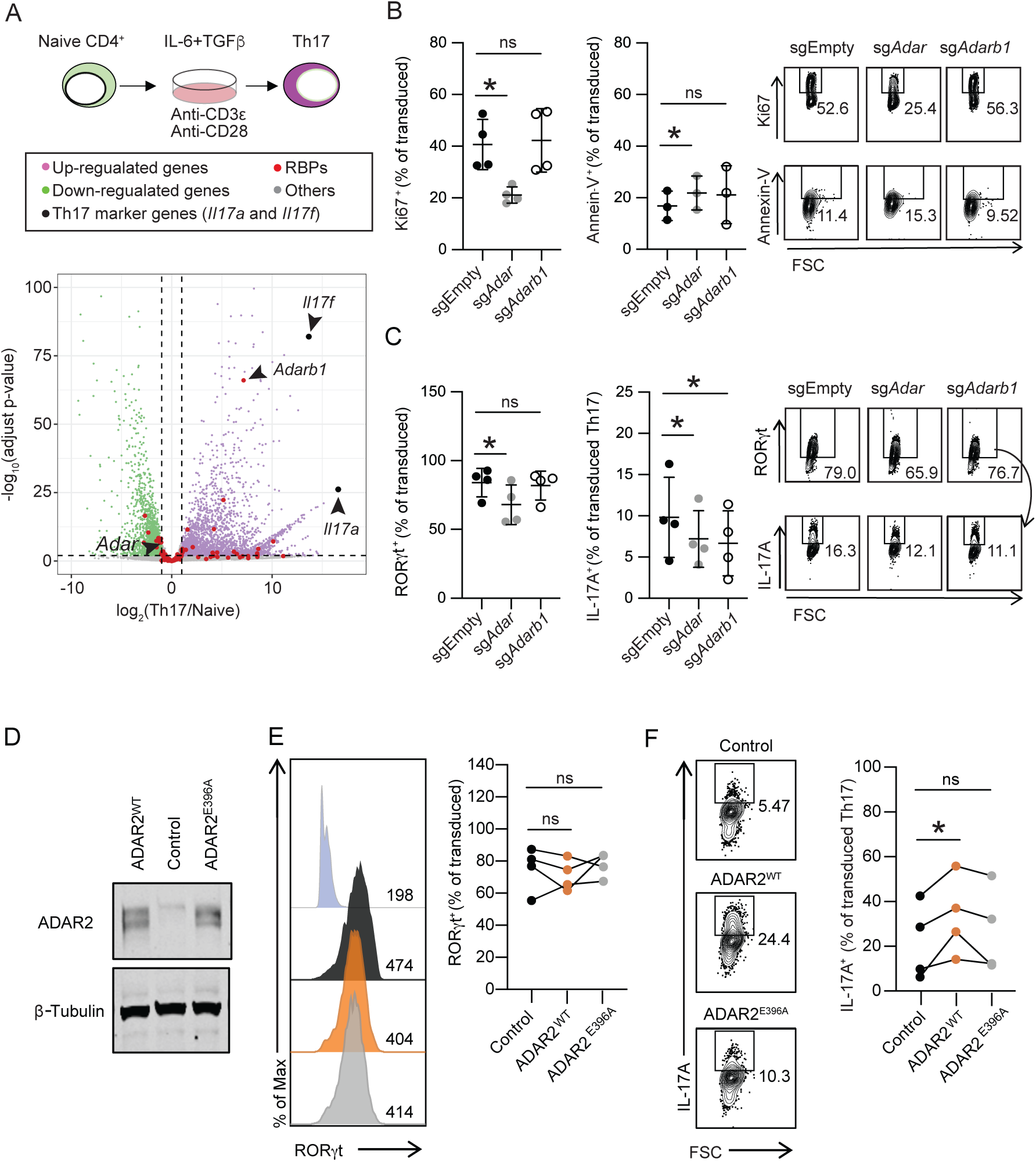
ADAR1 and ADAR2 regulate distinct aspects of Th17 biology. A. Cell culture assay workflow (top): purified murine naïve CD4^+^ T cells were activated and differentiated toward the Th17 lineage *in vitro*. Volcano plot (bottom) depicting differential gene expression in naïve and Th17 cells from two pairs of mice. Green and magenta dots indicate genes differentially expressed in naïve and Th17 cells (folds change >2 or <-2 and p<0.01). Grey dots indicate all other genes. Red dots indicate genes encoding RNA binding proteins (RBPs). *Il17a* and *Il17f* are highlighted for reference. B. Summarized proportions (left) and representative flow cytometry analysis (right) of Ki67 and Annexin-V in transduced (CD90.1^+^) Cas9^+^CD4^+^ T cells. FSC: Forward scatter as an indicator of cell size. * p-value<0.05, ns: not significant (paired t-test). C. Summarized proportions (left) and representative flow cytometry analysis (right) of RORγt in transduced CD90.1^+^CD4^+^ cells and IL-17A^+^ in transduced Th17 cells (defined as CD90.1^+^CD4^+^RORγt^+^). FSC: Forward scatter as an indicator of cell size. * p-value<0.05, ns: not significant (paired t-test). D. Representative western blot of whole-cell extracts from Th17 cells transduced with control, ADAR2^WT^, or enzymatically dead ADAR2^E396A^ expression constructs. This experiment was repeated twice on independent biological samples with similar results. E. Representative flow cytometry analysis (left) and summarized proportions (right) of RORγt^+^ transduced (CD90.1^+^CD4^+^) T cells. Numbers on the histogram indicate the mean fluorescent intensity of RORγt in each cell population. Isotype control (blue), empty vector transduced control (black), ADAR2^WT^ expression vector transduced (orange), ADAR2^E396A^ expression vector transduced (grey). Connecting lines indicate results from one independent experiment. ns: not significant (paired t-test, n=4). F. Representative flow cytometry analysis (left) and summarized proportions (right) of IL-17A^+^ Th17 cells in indicated transduced cells. Connecting lines indicate results from one independent experiment. * p-value<0.05, ns: not significant (paired t-test, n=4).

### ADAR1 and ADAR2 modulate distinct aspects of T cell biology

Next, we tested the function of ADAR1 and ADAR2 in an *in vitro* Th17 differentiation and effector function assay using the CRISPR-Cas9-mediated knockdown approach (Figure S1B). Depletion of ADAR1 significantly impaired cell proliferation as indicated by reduced Ki67-positive cells and increased cell apoptosis as evidenced by slight elevated Annexin-V-positive cells (Figure 1B). ADAR1-deficient cells also impaired the expression of the Th17 master transcription factor RORγt and effector cytokine IL-17A (Figure 1C). In contrast, depletion of ADAR2 did not alter cell proliferation, apoptosis, and RORγt expression levels, yet significantly lowered the production of IL-17A in Th17 cells (Figure 1B-C). These results suggest that while ADAR1 broadly modulates T cell survival, proliferation, and differentiation, ADAR2 serves a more specialized role in controlling Th17 effector function post differentiation.

### RNA editing activity of ADAR2 potentiates cytokine production in cultured Th17 cells

In cultured Th17 cells, overexpression of ADAR2^WT^ significantly increased IL-17A expression without altering the expression of RORγt (Figure 1D-F), consistent with the observation in the loss of function assay described above. Interestingly, cultured Th17 cells expressing the enzymatically dead ADAR2^E396A^ showed similar IL-17A production as control vector transduced cells (Figure 1F). This suggests that ADAR2 positively regulates Th17 effector function in an RNA editing-dependent manner.

To elucidate the RNA editomes in naïve and Th17 cells, we analyzed RNA-seq datasets from two biological replicates (Table S2-3) using a previously described workflow (*36*). Compared to Th17 cells, we detected more RNA variants in naïve T cells at both the site and gene levels (Figure 2A and S2A-C). 52.3% of base conversion variants identified in naïve CD4^+^ T cells and 62.9% in Th17 cells were A-to-G or T-to-C, indicative of A-to-I conversions (Figure S2A). The majority of A-to-I edited sites localized to 3’ untranslated regions (UTRs) (Figure S2D), in line with previous reports on other cell types (*37, 38*). Notably, however, *Il17a* transcripts did not harbor any A-to-I conversion in Th17 cells, suggesting that ADAR2 may modulate *Il17a* expression through a more elaborated mechanism.

**Figure 2.**
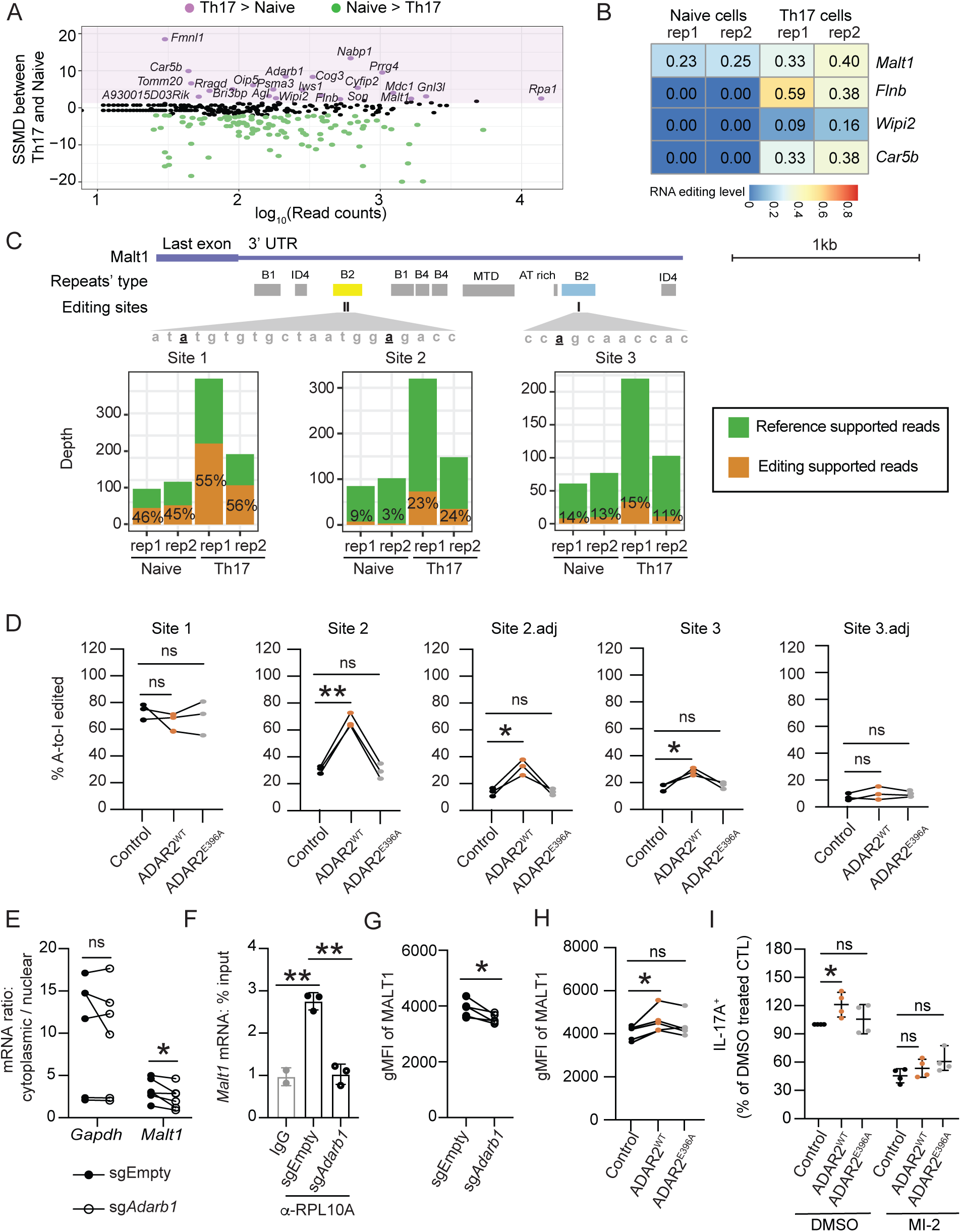
ADAR2 promotes IL-17A expression by editing MALT1. A. Strictly standardized mean difference (SSMD) of overall editing at the gene level between Th17 and naïve cells. Magenta and green dots indicate the genes differentially edited in Th17 and naïve cells (cut-off: SSMD>2 or SSMD<-2). Black dots indicate all other genes. B. Overall editing levels of the indicated genes in two independent replicates of naïve and Th17 cells. C. Position of repeat elements and editing sites on the *Malt1* 3’ UTR (top). Naïve and Th17 RNA sequencing depth and the fraction of reads containing A-to-I edits at each site (bottom). Vertical black lines indicate the editing sites identified by RNA-seq. D. Editing proportion of *Malt1* in Th17 cells transduced with the indicated vectors as determined by RT-PCR and Sanger sequencing. Connecting lines indicate results from one independent experiment. * p-value<0.05, ** p-value<0.01, ns: no significant (paired t-test, n=3). E. Ratio of *Malt1* mRNA detected in the cytoplasm and nucleus of transduced Cas9^+^ Th17 cells. Connecting lines indicate results from one independent experiment. * p-value<0.05, ns: not significant (paired t-test, n=6). F. Ribosome RPL10A enrichment on the *Malt1* transcript in transduced Cas9^+^ Th17 cells as detected by RIP and RT-qPCR. ** p-value<0.01 (t-test, n=3). G. Summarized geometric mean fluorescence intensity (gMFI) of intracellular MALT1 proteins in Cas9^+^ Th17 cells (CD90.1^+^CD4^+^RORγt^+^) transduced with the indicated sgRNA expressing vectors. Connecting lines indicate results from one independent experiment. * p-value<0.05 (paired t-test, n=5). H. Summarized geometric mean fluorescent intensity (gMFI) of intracellular MALT1 protein in Th17 cells transduced with the control or indicated ADAR2 expressing vectors. Connecting lines indicate results from one independent experiment. * p-value<0.05, ns: no significant (paired t-test, n=5). I. The fraction of IL-17A^+^ Th17 cells transduced with the indicated vectors normalized to those found in control vector transduced (CTL) cultures treated with DMSO. MI-2: MALT1 inhibitor, 500 nM. Each dot represents the results from one mouse. * p-value<0.05, ns: no significant (paired t-test, n=4).

### ADAR2 edits B2 SINE repeats on the *Malt1* 3′ UTR to promote translation

Fifteen transcripts harboring differential A-to-I editing in naïve vs Th17 cells encode proteins with altered abundance as quantified by mass spectrometry (Figure S3A) (*39*). Of the 10 proteins with greater expression in Th17 cells, 7 exhibited reduced A-to-I editing in their transcripts. In contrast, the degree of A-to-I editing on transcripts for ADAR2, MALT1, and SON positively correlated with their protein abundance in Th17 cells. These findings suggest that editing may modulate translation in a bidirectional fashion.

We were particularly intrigued by the extensive A-to-I editing on *Malt1* transcripts, as it represents one of the four transcripts that gained A-to-I editing in Th17 cells with a known role in defense response (Figure 2B and S3B). RNA-seq analysis revealed three major A-to-I edited sites (1–3) within two B2 SINE repeat elements on the *Malt1* 3’UTR (Figure 2C). Sanger sequencing of cDNAs from multiple independent Th17 cultures confirmed the presence of these three A-to-I conversion sites and identified two additional conversion sites nearby (2adj and 3adj) (Figure 2D). Native RNA immunoprecipitation (RIP) assay using an ADAR2-specific antibody confirmed significant enrichment of *Malt1* transcripts on ADAR2-containing complexes (Figure S3C). Overexpression of wildtype, but not mutant ADAR2, increased A-to-I editing on sites 2, 2adj, and 3 (Figure 2D and S4A), and conversely, ADAR2 knockdown reduced A-to-I editing on sites 2 and 2adj in cultured Th17 cells (Figure S4B-C). Notably, ADAR2 did not alter the overall *Malt1* mRNA output as indicated by the qRT-PCR analysis (Figure S5A) nor affect *Malt1* mRNA stability in Th17 cells treated with flavopiridol, a small molecular that globally inhibits mRNA transcription (*40*) (Figure S5B). Lastly, none of the *Malt1* A-to-I edited sites overlap with the TargetScan predicted microRNA target sites (Figure S5C). Collectively, these results suggest that ADAR2-mediated editing may directly impact MALT1 translation in Th17 cells.

Inverted repeats on mRNA 3’UTRs can form stem-loop structures that serve as binding sites for specific RBPs and regulate RNA localization to the nuclear paraspeckles and/or cytoplasmic stress granules (*41, 42*). Therefore, we speculated that ADAR2 editing of the inverted B2 SINE repeats on the *Malt1* transcript might alter RNA secondary structure, which may in turn affect its cellular localization and/or translation. In line with this possibility, the inverted B2 SINE repeats on *Malt1* were predicted to form a stem-loop-like structure and modifications of the adenosines at sites 2 and 2adj on the first B1 SINE repeat would disrupt the predicted structure (Figure S6A). In ADAR2 knockdown cultured Th17 cells, we saw a modest increase in nuclear retention of *Malt1* mRNA (Figure 2E). For the *Malt1* transcripts that made their way out to the cytoplasm, their engagement with the protein translation machinery was significantly reduced in the ADAR2-deficient cells (Figure 2F), consistent with the overall lowered MALT1 protein levels (Figure 2G). Conversely, MALT1 protein abundance was significantly elevated in Th17 cells overexpressing wildtype but not the enzymatic dead mutant of ADAR2 (Figure 2H). Together, these results demonstrate that ADAR2-mediated RNA editing promotes MALT1 expression at the nuclear export and protein synthesis steps in Th17 cells.

### The ADAR2-MALT1 axis regulates Th17 effector function *in vitro*

MALT1 is essential for IL-17A production in Th17 cells (*19, 20, 43, 44*). In line with these reports, we found that knockdown of *Malt1* or pharmacological inhibition of MALT1 protease activities significantly attenuated IL-17A production in culture Th17 cells (Figure S7A-D). Therefore, we hypothesized that ADAR2 may regulate IL-17A production and effector function of Th17 cells by regulating MALT1 levels. Consistent with this possibility, overexpression of ADAR2^WT^ in Th17 cells cultured in the presence of the MALT1 protease activity inhibitor (MI-2) failed to potentiate IL-17A production (Figure 2I). These results highlight that the ADAR2-MALT1 axis promotes IL-17A cytokine production in Th17 cells.

### ADAR2 protects against T cell-mediated colonic inflammation

In the intestine, Th17 cells enhance barrier function (*4–6*) and restrict local inflammation (*45*). Neutralization of IL-17A or its receptor worsens intestine inflammation in patients with inflammatory bowel disease (*11, 12*). Similar results were also observed in mice challenged with CD4^+^ T cell-transfer colitis (*4, 46*). Sanger analysis of the colonic lamina propria mononuclear cells in the CD4^+^ T cell-transfer colitis model confirmed abundant A-to-I editing on *Malt1* transcripts *in vivo* (Figure 3A). Therefore, we hypothesized that the ADAR2-MALT1 axis might also regulate Th17 cell effector function *in vivo* in settings of colitis. To test this possibility, control or ADAR2 overexpressing T cells were transferred into RAG1^-/-^ mice. By Day 35, recipients of ADAR2 overexpressing cells (ADAR2^WT^) were better protected from weight loss and had lower tissue pathology scores (Figure 3B-C). Th17 cells transduced with ADAR2-overexpressing constructs showed higher MALT1 protein levels than untransduced Th17 cells in the colonic lamina propria (cLP) and spleen of the same host (Figure 3D). In contrast, recipients of cells overexpressing the catalytic dead mutant (ADAR2^E396A^) showed similar weight changes compared to recipients of control cells (Figure 3E). These results demonstrate that ADAR2 augments MALT1 expressions in T cells *in vivo* and its deaminase activity protects against T cell-mediated colitis.

**Figure 3.**
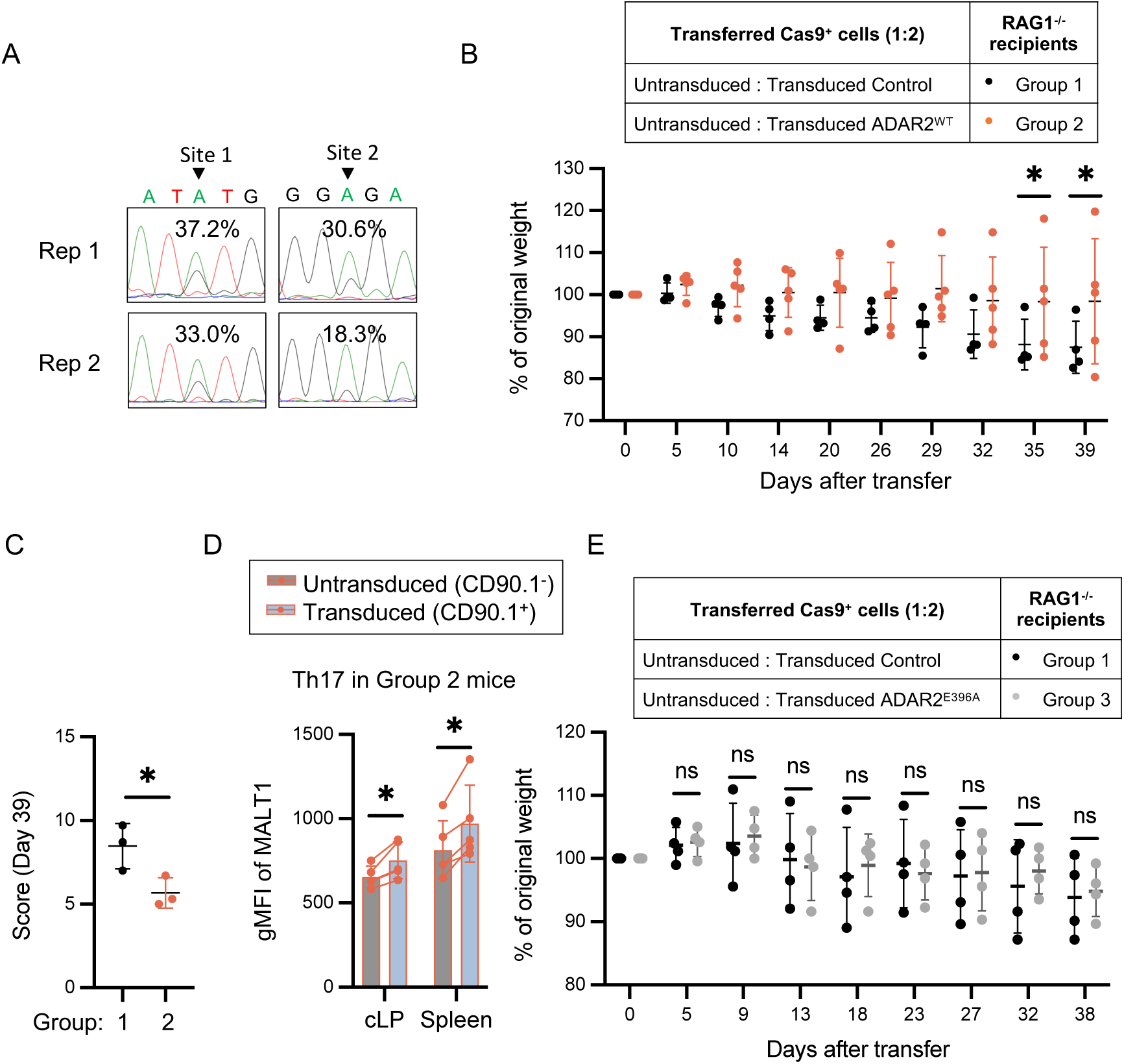
ADAR2 deaminase activity protects against T cell-mediated colitis. A. Representative Sanger sequencing analyses of A-to-I conversion at sites 1 and 2 of the *Malt1* mRNA in the colonic lamina propria (cLP) mononuclear cells. B. Weight change of RAG1^-/-^ mice receiving activated T cells transduced with empty (control) or ADAR2^WT^ overexpression constructs. Each dot represents the result from one mouse. * p-value<0.05 (t-test). C. Histology score of three pairs of colonic sections from B. * p-value<0.05 (paired t-test). Each dot represents the result from one mouse. D. The geometric mean fluorescent intensity (gMFI) of MALT1 in Th17 cells (CD4^+^ RORγt^+^) from the cLP and spleen of mice in group 2. Each connecting line represents the result from one mouse. * p-value<0.05 (t-test, n=5). E. Weight change of RAG1^-/-^ mice receiving activated T cells transduced with empty (control) or ADAR2^E396A^ overexpression constructs. Each dot represents the result from one mouse.

In a parallel experiment, activated CD4^+^ T cells from Cas9^+^ transgenic mice transduced with empty or sg*Adarb1* expression constructs were transferred into RAG1^-/-^ mice. Compared to the control, recipients of ADAR2-depleted (sg*Adarb1*) cells experienced greater weight loss by day 29 (Figure 4A). Colons from recipients of sg*Adarb1* cells harvested 35 days post-transfer displayed exacerbated pathology (Figure 4B-C) and had reduced overall tissue length (Figure 4D). A-to-I editing at site 2 of the *Malt1* transcripts displayed ADAR2 dependency *in vivo* (Figure 4E). Consistent with results from our cell culture experiments, ADAR2-depleted Th17 cells in the colonic lamina propria also showed lower IL-17A production capacity (Figure 4F). Interestingly, IFNγ, a cytokine produced by Th1 cells previously implicated in colitis, did not display ADAR2 dependency *in vivo* (Figure 4F). Furthermore, knocking down ADAR2 did not alter the proportions of colonic Th17, Th1, and Treg (Figure 4G, following the gating strategy on S8A). Importantly, these functional consequences recorded in ADAR2 deficient Th17 cells essentially phenocopy the functional defects of MALT1 knocked out Th17 cells (see Discussion), thus establishing the regulatory function of the ADAR2-MALT1 axis *in vivo*. Together, these results suggest that ADAR2 promotes Th17 effector function *in vivo* and protects against exacerbated colonic inflammation.

**Figure 4.**
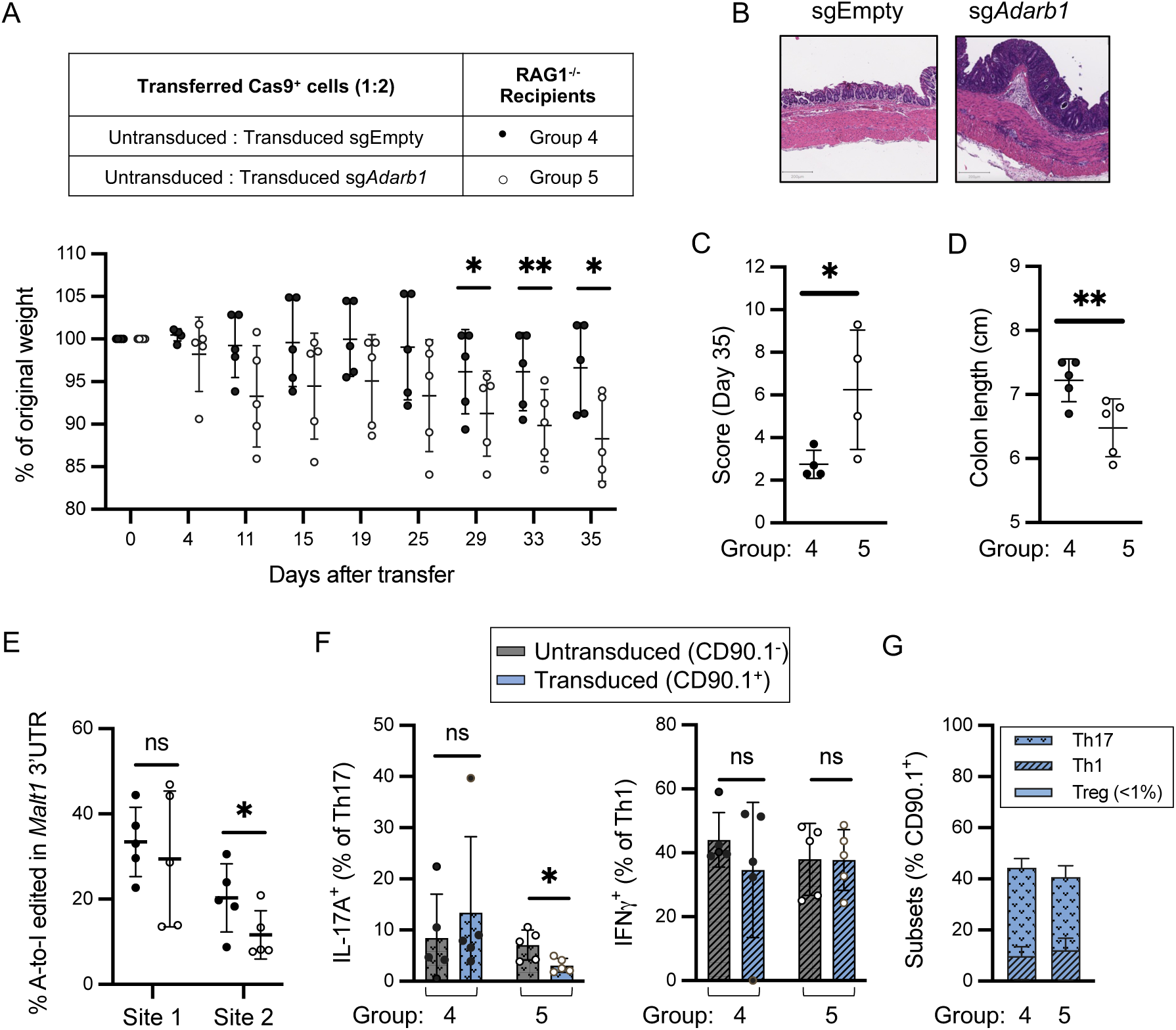
ADAR2 promotes Th17 effector function *in vivo*. A. Weight change of RAG1^-/-^ mice receiving activated Cas9^+^CD4^+^ T cells that were transduced with sgEmpty (group 1) or sg*Adarb1* (group 2). * p-value<0.05, ** p-value<0.01 (t-test, n=5). Each dot represents the result from one mouse. The results shown are combined from two independent experiments. B. Representative H&E staining of colonic sections harvested on day 35 after transfer. Scale bar: 200μm. C. Summarized histology score of B. Each dot represents the result from one mouse. * p-value<0.05 (t-test). D. Colon length of the mice from A harvested on day 35 after transfer. Each dot represents the result from one mouse. ** p-value<0.01 (t-test, n=5). E. The percentage of A-to-I edited Malt1 transcripts in the cLP mononuclear cells. * p-value<0.05, ns: no significant (t-test). F. The proportion of IL-17A^+^ Th17 and IFN*γ*^+^ Th1 cells in cLP. Each dot represents the result from one mouse. * p-value<0.05, ns: no significant (t-test). G. The proportion of Th17 (RORγt^+^CD4^+^), Th1 (Tbet^+^CD4^+^) and Treg (Foxp3^+^CD4^+^) cells in cLP.

### Super enhancer promotes *Adarb1* transcription during Th17 polarization

Differentiation of naïve CD4^+^ T cells toward different T helper lineages requires drastic changes to the chromatin landscape. Most notable is the emergence of super-enhancers encompassing large clusters of enhancers enriched with the active H3K27 acetylation (H3K27ac) marks (*47–50*). To elucidate the mechanism underlying the induction of ADAR2 expression in Th17 cells (Figures 1A and 5A), we characterized the chromatin landscape of the *Adarb1* locus using the Chromatin Immunoprecipitation with high-throughput sequencing (ChIP-seq) and the Assay for Transposase-Accessible Chromatin with high-throughput sequencing (ATAC-seq). As shown in Figure 5B-C, the *Adarb1* locus underwent dynamic opening when naïve cells were polarized toward Th17. 72hrs post differentiation into Th17, genome-wide characterization of histone H3 lysine 27 acetylation (H3K27ac) revealed the establishment of a putative intragenic super-enhancer (SE) at the *Adarb1* locus (Figure 5B). This SE spanned a cluster of five cis non-coding sites (CNS) that gained accessibility during Th17 polarization (CNS1: chr10 77367402-77367422, CNS2: chr10 77368829-77368848, CNS3: chr10 77387874-77387894, CNS4: chr10 77388804-77388824, and CNS5: chr10 77406713-77406733) (Figure 5C). To further understand how this SE interacts with the *Adarb1* promoter, we performed Global RNA interactions with DNA by deep sequencing (GRID-seq, (*51, 52*)) to obtain evidence for the proximity of the SE with the nascent *Adarb1* RNA. Indeed, we found that numerous cis elements within this putative SE were positioned in close spatial proximity to the *Adarb1* promoter (Figure 5C). Analysis of publicly available datasets (*53, 54*) further revealed that CNS 3 and 4 occupied opened chromatins only in Th17 cells and not any other CD4^+^ T cell subsets (Figure S9A-B).

**Figure 5.**
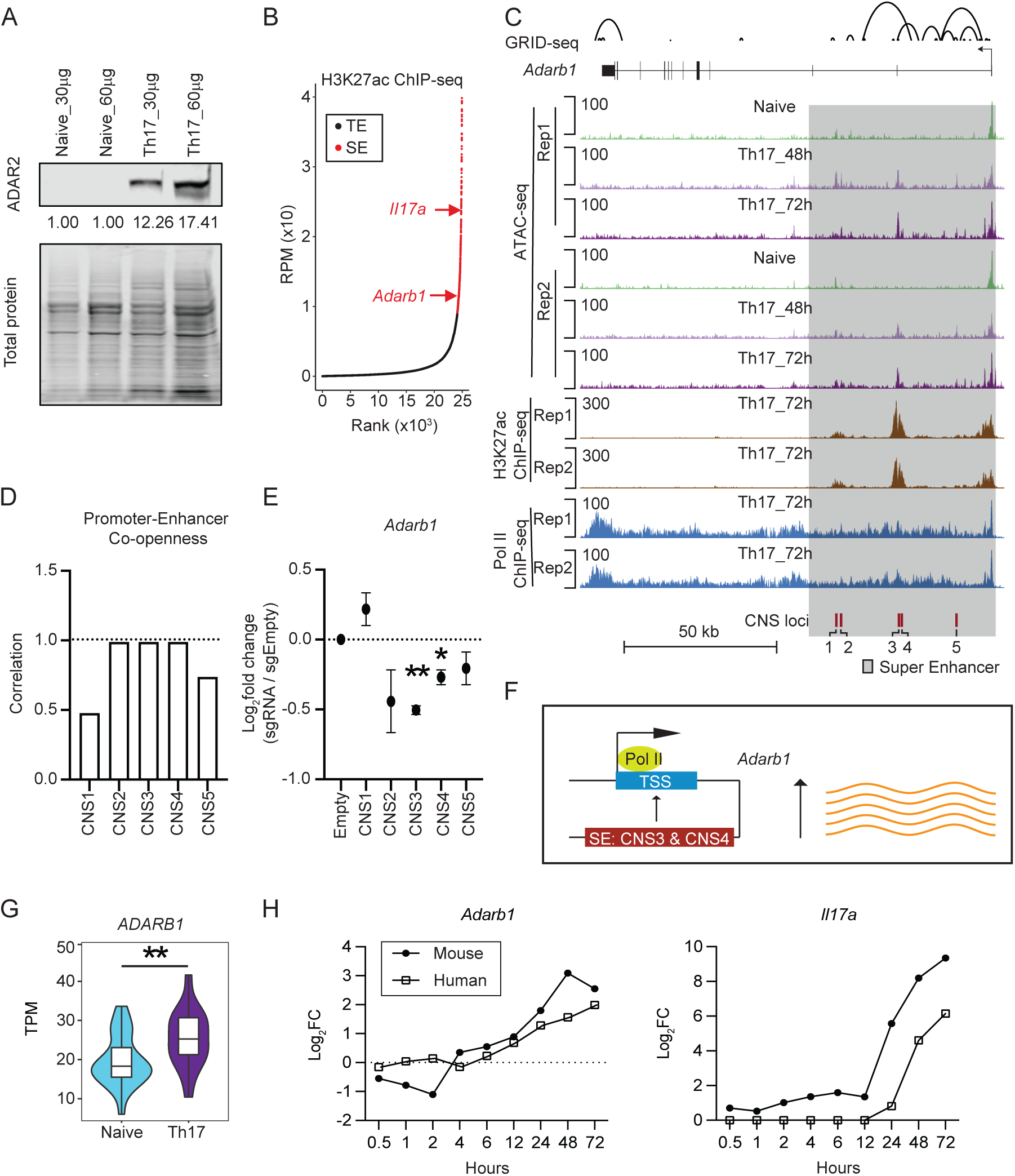
*Adarb1* transcription is controlled by an intragenic super-enhancer in Th17 cells. A. Representative western blot analysis of whole-cell extracts from naïve CD4^+^ T and cultured Th17 cells harvested 72hrs post differentiation. This experiment was repeated twice on independent biological samples with similar results. B. Th17 super-enhancers (SE, red dots) and typical enhancers (black dots) as defined by regional H3K27ac signals (see Methods for detail). The arrow highlights the SE of *Adarb1*. C. Chromatin landscape of the mouse *Adarb1* locus. Chromatin accessibility assays were performed on two independents naïve CD4^+^ T and cultures of Th17 cells harvested at 48 or 72 hours. ChIP-seq (acetylated H3K27 and RNA Polymerase II) and GRID-seq were performed on two independent cultures of Th17 cells harvested at 72 hours. Shaded box: putative super-enhancer region. D. Co-openness correlation scores of the *Adarb1* promoter and each putative cis non-coding site (CNS). E. Log_2_ fold changes in *Adarb1* mRNA quantified by qRT-PCR in Cas9^+^ Th17 cells transduced with sgRNAs targeting the corresponding CNS compared to control. Average and standard deviation of results from two independent experiments are shown. * *p*-value<0.05, ** *p*-value<0.01 (t-test). F. Working model: CNS3 and CNS4 within the *Adarb1* intragenic super-enhancer promote local transcription in murine Th17 cells. G. Violin plot of human *ADARB1* expression in naïve and cultured Th17 cells from 5 independent RNA-seq datasets (*43*). H. Log_2_ fold change of *ADARB1* and *IL17A* expression in murine or human Th17 cells at different time points compared to naïve controls as determined by RNA-seq (*44*).

Among these five CNSs, chromatin opening at CNS2, CNS3, and CNS4 exhibited the highest correlation with the accessibility dynamics at the *Adarb1* promoter, suggesting that these elements may regulate transcription of the *Adarb1* locus (Figure 5D). To assess their contribution(s) to *Adarb1* expression, small guide RNAs (sgRNAs) were designed to target each of the putative elements in Cas9-expressing Th17 cells. CRISPR-mediated mutation of CNS3 or CNS4 resulted in a significant reduction in *Adarb1* expression (Figure 5E). Altogether, these results revealed that CNS3 and CNS4 within the *Adarb1* SE are essential for driving ADAR2 expression in Th17 cells (modeled in Figure 5F).

Induction of ADAR2 is conserved in human Th17 cells. Analysis of ten public human naïve and/or Th17 datasets (*55*) by ComBat-seq under default parameters revealed that *ADARB1* was also significantly upregulated when human naïve CD4^+^ T cells were polarized toward the Th17 lineage (Figure 5G and S10A-B). With an independent time-course dataset (*56*), we further confirmed the striking parallel induction of *ADARB1* and *IL-17A* transcripts during human and murine Th17 cell polarization (Figure 5H), suggesting an evolutionarily conserved association between ADAR2 and acquisition of Th17 effector function.

### The ADAR2-MALT1 axis is conserved in human

Analysis of the public RNA-seq datasets from human Th17 and non-Th17 cells revealed the presence of extensively edited Alu repeats on the human *MALT1* 3’UTR (Figure 6A). To test the possibility that human ADAR2 also serves as the upstream regulator of MALT1, we over-expressed two isoforms of ADAR2 most abundantly expressed in human T cells in HEK293 cells (Figure 6B). A previous study reports that the *ADARB1*(+5a) transcript containing the alternatively spliced exon 5a (ENST00000437626.5) resulting in the insertion of an AluJ cassette in the ADAR2 catalytic domain, encodes a less catalytically active enzyme compared to the one encoded by the shorter *ADARB1* (ENST00000348831.9) transcript (*57*). As expected, over-expression of *ADARB1* and, to a lesser extent, *ADARB1*(+5a) increased A-to-I editing on select sites on *MALT1* 3’UTR (Figure 6C and S11A) and potentiated MALT1 protein expression modestly by 20-30% (Figure 6D). Notably, overexpression of ADAR2 did not alter ADAR1 protein levels in HEK293 cells (Figure 6D). Altogether, these data suggest that ADAR2 regulation of *MALT1* is an evolutionarily conserved pathway that can be utilized by both immune and non-immune cells.

**Figure 6.**
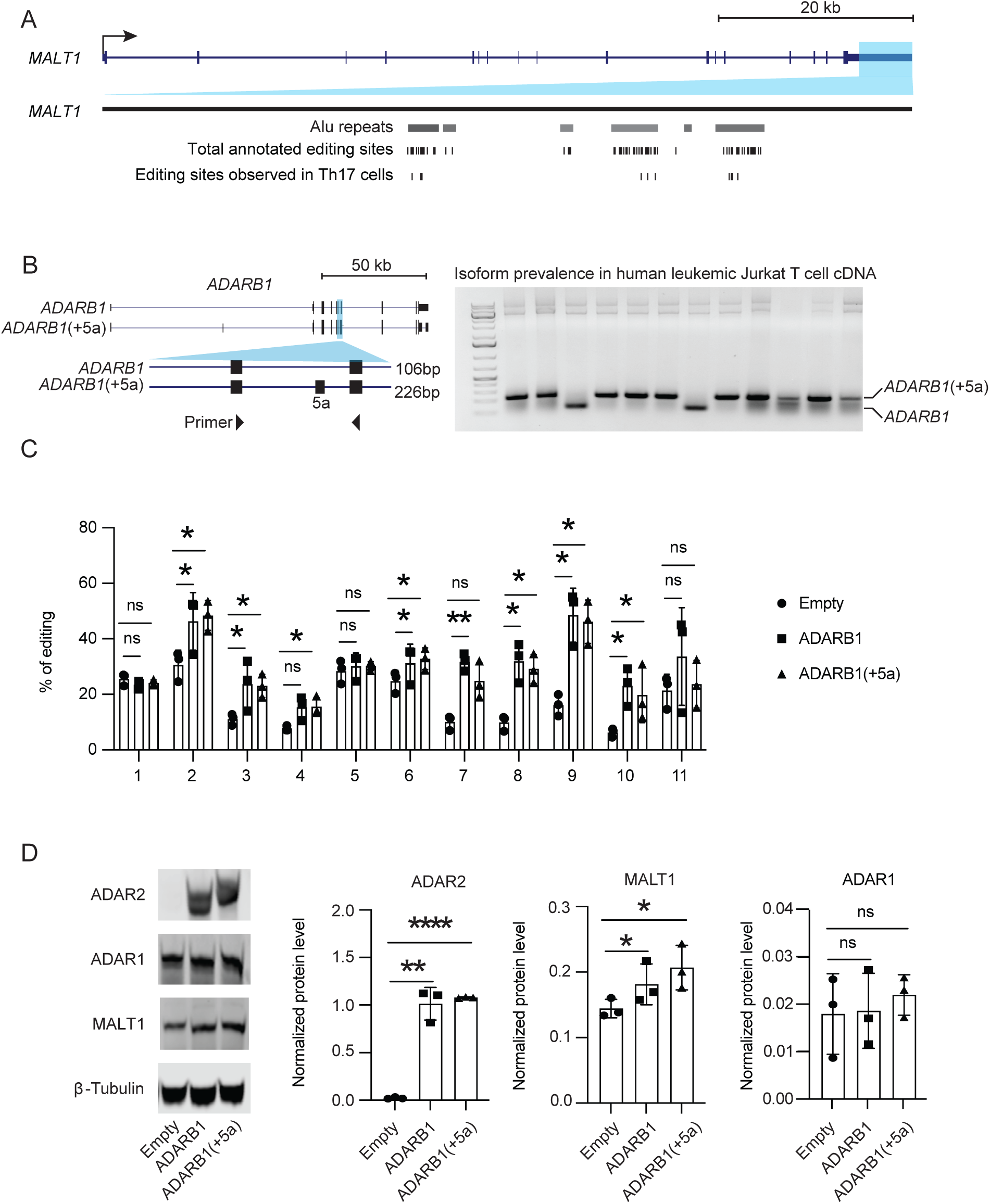
The ADAR2-MALT1 pathway is conserved in humans. A. Location of Alu repeats and A-to-I editing sites on the *MALT1* 3’ UTR in human Th17 and non-Th17 cells. B. Schematic of two *ADARB1* isoforms (left), *ADARB1* and ADARB1(+5a), expressed in Jurkat T cells. PCR results (right) using the indicated primers on 12 independent cDNA clones obtained from Jurkat T cell. C. The proportion of A-to-I conversion on *MALT1* in HEK293 cells transfected with the indicated vectors was determined by RT-PCR and Sanger sequencing. Connecting lines indicate results from one independent experiment. * p-value<0.05, ** p-value<0.01, ns: no significant (paired t-test, n=3). D. Representative (left) and quantification of the indicated proteins in control or HEK293 cells over-expressing the indicated *ADARB1* isoform as determined by western blot analysis. β-tubulin: loading control used for normalization. Connecting lines indicate results from one independent experiment. * p-value<0.05, ** p-value<0.01, **** p-value<0.0001, ns: no significant (paired t-test, n=3).

## DISCUSSION

Prior studies on the molecular regulation of T cell differentiation and function primarily focused on transcription factors, but the role of RNA binding proteins (RBPs) in these processes remains poorly understood. Here, we identify ADAR2 as an important regulator of Th17 effector function. Notably, ADAR2 is the most highly upregulated RBP when naïve helper T cells differentiate toward the Th17 lineage. Results from our genetic and genomic experiments demonstrate that the hyperactivation of the *Adarb1* locus in Th17 cells is driven by a novel intragenic super-enhancer. Once induced, ADAR2 promotes the production of Th17 effector cytokine, IL-17A, *in vitro* and *in vivo* in a deaminase activity dependent manner. This regulation is, in part, facilitated by ADAR2-mediated editing on MALT1 transcripts. Genetic ablation of *Malt1* during thymic T cell development and/or pharmacologic inhibition of MALT1 proteolytic activity result in multi-organ inflammation (*44, 58–62*). In Th17 cells, MALT1 inactivates RelB, promotes p65 nuclear translocation, and drives the expression of IL-17A (*19, 20, 43, 44*). In models of neuro-inflammation, MALT1 regulates the expression of GM-CSF and IL-23 and promotes pathogenic Th17 functions (*19, 20, 58, 63*). In this study, we show that depletion of ADAR2 dampens MALT1 expression and reduces the capacity of Th17 cells to produce IL-17A *in vitro. In vivo,* it is possible that MALT1 together with other molecules that are also regulated by RNA editing contribute to the protective role of ADAR2 during colitis.

Contrary to the previous reports suggesting that ADAR2 primarily catalyzes adenosine deamination of non-repetitive RNA sequences while ADAR1 edits repetitive elements (*21*), ADAR2 modifies select adenosines within two inverted B2 SINE repeats on the murine *Malt1* transcript and the Alu repeats on the human *MALT1* transcript. SINE and Alu repeats are the most abundant repetitive elements in the mouse and human genomes, respectively. An estimated 4% of these repeats localize to the 3’ UTRs of fully spliced transcripts (*64*). Inverted repeats on 3’ UTRs restrict protein synthesis by retaining transcripts in nuclear paraspeckles and/or cytoplasmic stress granules (*41, 42*). But the regulatory molecule(s) involved in releasing target transcripts from this type of negative regulation is not well understood. Here, we uncover ADAR2 mediated A-to-I editing as an evolutionarily conserved mechanism for unleashing the translation potential of invert repeats-bearing transcripts. By editing the *Malt1* 3’ UTR inverted B2 SINEs, ADAR2 facilitates *Malt1* export to the cytoplasm and promotes translation.

In addition to ADAR2, other adenosines on the *Malt1* transcript are targeted by ADAR1. While ADAR2 exhibits tissue and T cell subset-specific expression, ADAR1 and MALT1 are ubiquitously expressed across immune and non-immune cells. In Th17 cells, however, the abundant expression of ADAR1 did not compensate for the loss of ADAR2, suggesting the two ADARs likely have non-redundant roles in regulating the *Malt1* transcript. To investigate whether the two enzymes contribute to MALT1 expression and other aspects of Th17 function in cooperative or competitive manners, future studies will need to first bypass the essential role of ADAR1 in T cell survival and proliferation, possibly by generating an inducible system for depleting ADAR1 after Th17 commitment or in transgenic backgrounds with elevated expression of anti-apoptotic factors.

In contrast to the well-characterized transcriptional mechanisms involved in T cell development and differentiation, post-transcriptional switches exemplified by ADARs and RNA editing uncovered in this study provide a more rapid pathway for effector T cells to respond to challenges during tissue injury, inflammation, and repair. In summary, our study highlights the essential role of ADAR2 in regulating Th17 effector function and provides new molecular insights into the role of RNA editing in T cell and mucosal biology.

## ACKNOWLEDGEMENTS

We thank Johannes Zuber and Aichinger Martin at the Research Institute of Molecular Pathology (I.M.P.) for sharing the pSIN vector for generating retroviruses to carry custom sgRNAs into T cells.

S.M., B.S.C., N.C., A.Z., P.R.P., and W.J.M.H. are partially funded by the Edward Mallinckrodt, Jr. Foundation and the National Institutes of Health (NIH) (R01 GM124494 to WJM Huang). Y.H. was supported by NIH grants HG004659 and DK098808. Illumina sequencing was conducted at the IGM Genomics Center, University of California San Diego, with support from NIH (S10 OD026929) and Moores Cancer Center (P30 CA023100).

## AUTHOR CONTRIBUTIONS

S.M. designed and performed most *in vivo* and *in vitro* studies with some help from B.S.C., N.C and A.Z. Y.H performed the analyses on the RNA-seq datasets, editomes, and the Malt1 RNA structure with some help from S.Z. N.C. and B.S.C. completed the SANGER validation of the SAILOR analysis in Th17 cells. A.Z performed the human editing. A.Z. and P.R.P. completed the blinded histology analysis on colonic sections. B.Z. performed the enhancer and super-enhancer analyses on the ChIP-seq datasets. X.D.F. directed the analyses by Y.H, S.Z, and B.Z and edited the manuscript. W.J.M.H. analyzed and wrote the manuscript together with S.M.

## METHODS

### Mice

C57BL/6 wild-type (Stock No: 000664), Cas9^Tg/+^ (Stock No: 026179), and RAG1^-/-^ (Stock No: 002216) were obtained from the Jackson Laboratory. Adult mice, at least eight weeks old, were used. All animal studies were approved and followed the Institutional Animal Care and Use Guidelines of the University of California San Diego.

### Cell Culture

Mouse naive T cells were purified from spleens and lymph nodes of 8-12 weeks old mice using the Naive CD4^+^ T Cell Isolation Kit according to the manufacturer’s instructions (Miltenyi Biotec). Cells were cultured in Iscove’s Modified Dulbecco’s Medium (IMDM, Sigma Aldrich) and supplemented with 10% heat-inactivated FBS (Peak Serum), 50 U penicillin-streptomycin (Life Technologies), 2 mM glutamine (Life Technologies), and 50 μM β-mercaptoethanol (Sigma Aldrich). For Th17 cell polarization, naive cells were seeded in 24-well or 96-well plates pre-coated with rabbit anti-hamster IgG and cultured in the presence of 0.25 μg/mL anti-CD3ε (eBioscience), 1 μg/mL anti-CD28 (eBioscience), 0.1-0.3 ng/mL TGF-β (R&D Systems), and 20 ng/mL IL-6 (R&D Systems) for 48 to 72 hours. MALT1 inhibitor MI-2 (MedChemExpress, 50-1000 nM) or DMSO control were added together with the polarization cytokines.

The HEK293 cells were maintained in DMEM medium (Gibco) supplemented with 10% FBS and 50 U penicillin-streptomycin. Human *ADARB1* and *ADARB1*(+5a) were amplified from Jurkat T cell cDNAs and cloned into pcDNA3 (Invitrogen). HEK293 cells were transfected with the control and ADARB1 encoding pcDNA3 constructs using Lipofectamine 3000 (Invitrogen) and analyzed 72hrs post-transfection.

### Retrovirus Transduction in T cells

sgRNAs were designed by CHOPCHOP (*65*). pSIN retroviral constructs carrying expression cassettes for sgRNA along with a surface protein CD90.1 (Thy1.1) were generated as described previously in (*66*). sgRNA sequences are listed in Table S4. The MSCV wildtype murine ADAR2 overexpression construct was generated using the In-Fusion HD Cloning Plus kit (Takara Bio). Specifically, primers were designed using the Takara Primer Design tool with NotI and SalI extensions. The amplified PCR products were ligated to the NotI and SalI linearized MSCV vector (RRID: Addgene_17442). The MSCV murine ADAR2^E396A^ expression construct was generated using the Q5 Site-Directed Mutagenesis Kit (New England Biolabs). Clones were screened and confirmed by Sanger sequencing.

Retroviruses were generated in PlatE cells (*67*). Cas9^+^ murine naive CD4^+^ T cells were purified from spleens and lymph nodes of 8-12 weeks-old mice. Virus transduction on Cas9^+^CD4^+^T cells was performed 24 hours after T cell activation by centrifugation at 2000 rpm for 90 min at 32°C. Live and CD90.1^+^ retrovirus transduced cells were polarized and analyzed by flow cytometry at 72 hrs.

### T cell transfer colitis

For the ADAR2 loss of function study, 2 million activated Cas9^+^CD4^+^ T cells transfected with sgEmpty or sg*Adarb1* were injected intraperitoneally into RAG1^-/-^ recipients. For the ADAR2 gain of function study, 0.25 million sorted Cas9^+^CD4^+^CD90.1^+^ T cells transduced with MSCV-empty (control) or MSCV-ADAR2 (ADAR2^WT^) were injected intraperitoneally into RAG1^-/-^ recipients. Mice weights were measured twice a week and tissues were harvested between 35-39 days. Distal colons were collected for H&E staining. Pathology scoring of challenged mice was performed blind following previously published guidelines (*68*). Isolation of colonic lamina propria mononuclear cells was performed as previously described (*69*).

### Flow Cytometry

Cells were stimulated with 5 ng/mL Phorbol 12-myristate 13-acetate (PMA, Millipore Sigma) and 500 ng/mL ionomycin (Millipore Sigma) in the presence of GolgiStop (BD Bioscience) for 5 hours at 37 °C, followed by cell surface marker staining for CD4 (clone GK1.5; Biolegend) and CD90.1/Thy1.1 (clone HIS51; eBioscience). Fixation/Permeabilization buffers (eBioscience) were used as per manufacturer instructions to assess intracellular transcription factors and cytokine expression. Antibodies are listed in Table S5.

### RNA-seq

Ribosome-depleted RNAs from two independent replicates of naïve and culture Th17 cells were used to generate sequencing libraries. 100 bp paired-end sequencing was performed on an Illumina HiSeq4000 by the Institute of Genomic Medicine at the University of California San Diego. Each sample yielded approximately 30-40 million reads. Paired-end reads were aligned to the mouse mm10 genome with the STAR aligner version 2.6.1a (*70*) using the parameters: “--outFilterMultimapNmax 20 -- alignSJoverhangMin 8 --alignSJDBoverhangMin 1 --outFilterMismatchNmax 999 -- outFilterMismatchNoverReadLmax 0.04 --alignIntronMin 20 --alignIntronMax 1000000 -- alignMatesGapMax 1000000”. Uniquely mapped reads overlapping with exons were counted using featureCounts (*71*) for each gene in the GENCODE.vM19 annotation. Differential expression analysis was performed using DESeq2 (v1.18.1 package) (*72*), including a covariate in the design matrix to account for differences in harvest batch/time points. Regularized logarithm (rlog) transformation of the read counts of each gene was carried out using DESeq2. Differentially expressed protein-coding genes with minimal counts of 10, log2 fold change cutoffs of ≥ 0.5 or ≤ 0.5, and p-values <0.05 were considered significant.

For RNA variant analysis, low-quality reads were filtered out and adapters removed using Trimmomatic with parameters SLIDINGWINDOW:2:20 MINLEN:38. Ribosome RNAs reads were removed using SortMeRNA. RNA variations were called by the variants on RNA-seq data pipeline from the Genome Analysis Toolkit (GATK) following best practices described previously (*73*). Variants must be supported by at least one mismatched read with a base quality score ≥25, a mapping quality score ≥20, and coverage of each site ≥10. Only variations matching those previously curated by the REDIportal or DARNED database were included in the analysis. Gene-level editing scores were calculated as the number of edited reads mapping to the transcript over the total number of reads (edited and reference) mapping to the transcript (Figure S2C).

### ChIP-seq and ATAC-seq

ChIP-seq experiments were performed on 5-10 million Th17 cells crosslinked with 1% formaldehyde. Chromatin was sonicated and immunoprecipitated using antibodies listed in Table S5 and Dynabeads (Thermo Fisher Scientific), followed by reverse cross-linking, and library construction. ATAC-seq libraries were generated as described in (*74*). ChIP-seq and ATAC-seq processing followed the ENCODE guideline with some modifications. Specifically, single-end raw reads were mapped to the mouse genome (GENCODE assembly GRCm38) by bowtie2 (Version 2.3.4.1) in the local mapping mode with parameter “--local”, followed by PCR deduplication by SAMTools (Version 1.9) with the utility markedup (*75*). Mapped reads from the two replicates were merged into a single BAM file by SAMTools, and peaks were called using MACS2 (Version 2.2.6) (*76*) in the narrow peak-calling mode with default parameters for ChIP-seq data or specific parameters of “callpeak --nomodel --extsize 100” for ATAC-seq data. Regions with peak-score below 30 were filtered out and the remaining reliable peak profiles were transformed into bigwig format and visualized on the Integrative Genomics Viewer (IGV Version 2.8.2) (*77*).

### Identification of Enhancers and Super-enhancers

To identify super-enhancers, H3K27ac peak regions within a 12.5 Kb window outside of any known-gene promoters (TSS ± 2.5 Kb), were stitched together and ranked by increasing total background subtracted by ChIP-seq occupancy of H3K27ac. Ranked regions were plotted against the total background subtracted by ChIP-seq occupancy of H3K27ac in units of total rpm on the y-axis. Regions above the geometrically defined inflection point from this representation were considered putative super-enhancers.

Co-openness correlation scores of the *Adarb1* promoter and each putative cis non-coding site (CNS) were calculated using ATAC-seq signals from naïve, 48hrs, and 72hrs post polarization toward Th17 as previously described (*78*). The *Adarb1* promoter (Chr10: 77415285-77416735) with a Pearson correlation coefficient larger than 0.9 was selected as the potential target of the enhancers.

### Western Blot Analysis

For whole cell lysates, cells were lysed in 25 mM Tris pH 8.0, 100 mM NaCl, and 0.5% NP40 with protease inhibitors for 30 min on ice. Samples were spun down at 14,000 x g for 15 min, and soluble protein lysates were harvested. 20-50 μg proteins were loaded on each lane. Blots were blocked with the Odyssey Blocking buffer (Li-CoR Biosciences) and probed with primary antibodies listed in Table S5. Following incubation with respective IRDye secondary antibodies (Li-CoR Biosciences), infrared signals on each blot were captured by the Li-CoR Odyssey CLX.

### cDNA Synthesis, qRT-PCR, and RT-PCR Sanger Analysis

Total RNA was extracted with the RNeasy kit (QIAGEN) and reverse transcribed using SuperScript™ III First-Strand Synthesis System (Thermo Fisher Scientific). Real-time RT-PCR was performed using iTaq™ Universal SYBR® Green Supermix (Bio-Rad Laboratories). Expression data were normalized to *Gapdh* mRNA levels. qRT-PCR primers were designed using Primer-BLAST to span across splice junctions, resulting in PCR amplicons that span at least one intron. Primer sequences are listed in Table S4.

To assess the fraction of candidate transcripts carrying A-to-I edits, RT-PCR was performed on cDNAs from murine Th17 or human HEK293 cells using the Q5 Hot Start High-Fidelity 2X Master Mix (New England Biolabs) with the standard thermocycling protocol. Samples were incubated at 98 °C for 30 seconds, followed by a denaturation for 10 s at 98 °C, 20 s at 66 °C, and 72 °C for 1 min for a total of 34 cycles. PCR products were purified using Quick PCR Purification Kit (Qiagen) and sequenced. Primer sequences are listed in Table S4.

### RNA immunoprecipitation (RIP)

For RIP experiments, Th17 cells were lysed in 25 mM Tris pH 8.0, 100 mM NaCl, 0.5% NP40, 10 mM MgCl2, 10% glycerol with RNase and protease inhibitors for 30 min on ice. Samples were spun down at 14,000 x g for 15 min, and soluble protein lysates were harvested and immunoprecipitated with isotype control, ADAR2, or RPL10A-specific antibodies as listed in Table S5.

### Statistical Analysis

All values are presented as means ± SD. Significant differences were evaluated using GraphPad Prism 8 software. The student’s t-tests or paired t-tests were used to determine significant differences between two groups. A two-tailed p-value of <0.05 was considered statistically significant in all experiments.

Table S1. DESeq analysis of RBP genes in CD4^+^ naive and Th17 cells

Table S2. Editing site analysis of CD4^+^ naive and Th17 cells

Table S3. Editing level of genes in CD4^+^ naive and Th17 cells

Table S4. Primers and sgRNAs.

Table S5. Antibodies.

## SUPPLEMENTAL INFORMATION

**Figure S1.**
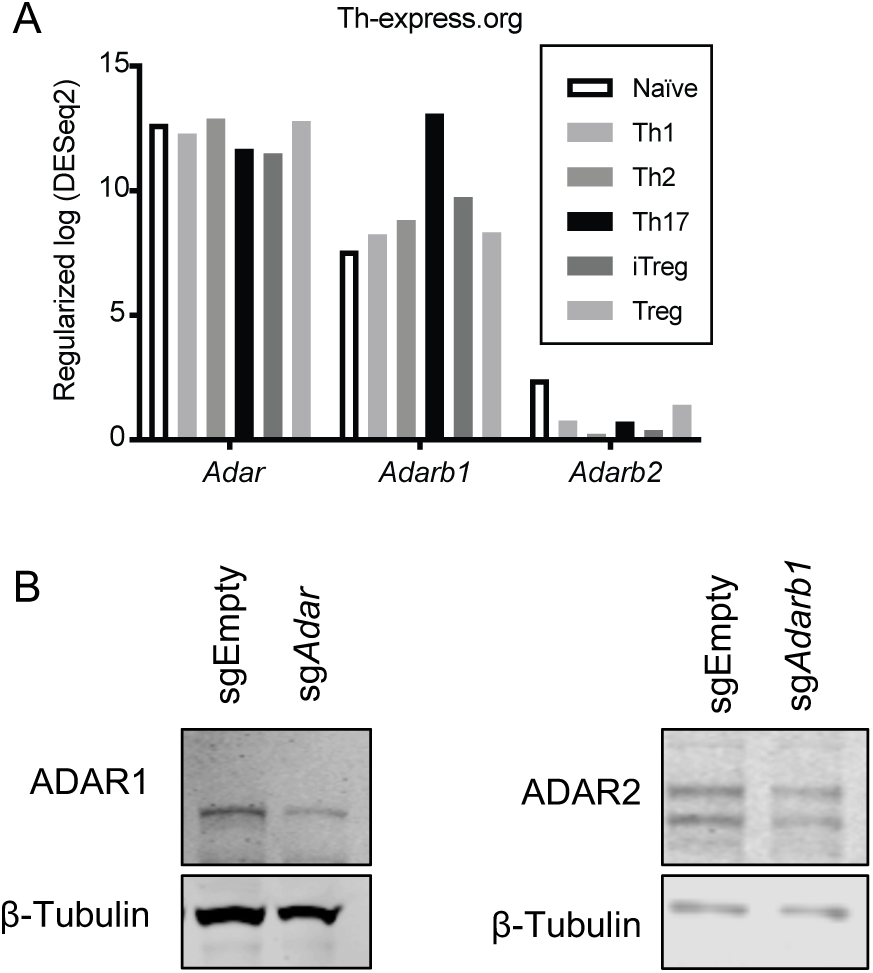
ADAR1 and ADAR2 expression in different T cell subsets. A. Regularized logarithmic transformed read counts of the indicated gene determined by RNA-seq (*29*). B. Representative western blot analysis of ADAR1 or ADAR2 in Th17 cells transduced with the indicated sgRNA expression constructs. This experiment was repeated twice on independent biological samples with similar results.

**Figure S2.**
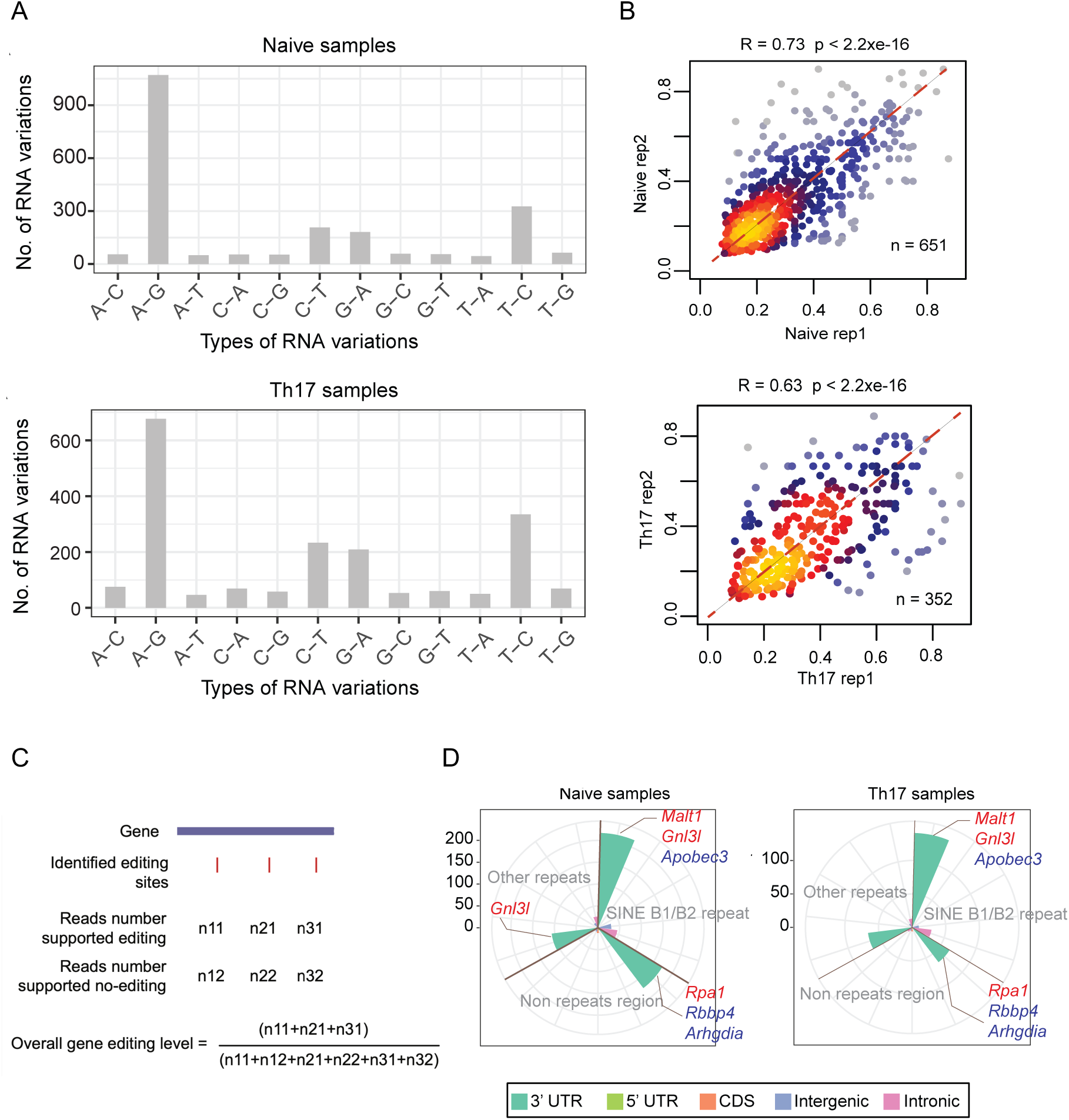
RNA editomes in T cells. A. Distribution of 12 types of RNA editing events in naïve (top) and culture Th17 cells (bottom). Data represent the total number of RNA editing events detected in both of replicates of the corresponding cell types. B. Scatterplot of A-to-I editing events identified in A that have been previously curated by the REDIportal or DARNED database on each independent biological replicates. Each dot represents one editing site. C. Workflow for calculating overall editing level at the gene level used for Figure 2A. D. Distribution of edited sites on different transcript regions in naïve and culture Th17 cells.

**Figure S3.**
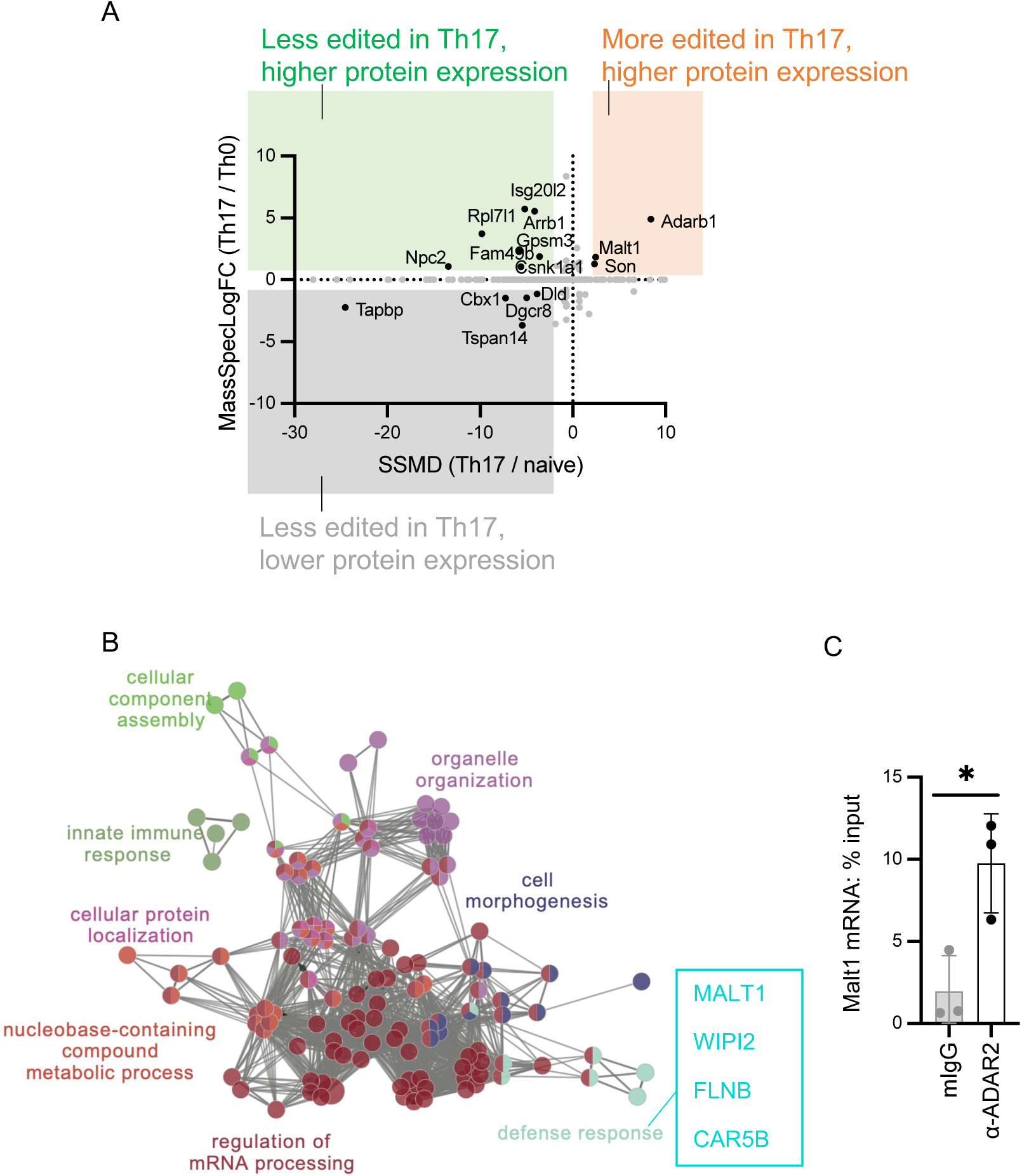
Protein abundance of A-to-I edited transcripts in naïve and Th17 cells. A. Scatterplot of protein abundance differences in Th17 and non-polarized Th0 (*33*) and transcript editing level changes (SSMD) in Th17 and naïve cells. Black dots: 15 transcripts with SSMD>2 or <-2 with differential protein abundance. Grey dots: all other transcripts. B. Gene ontology analysis of the 23 genes harboring higher editing in culture Th17 cells compared to naïve cells. Each dot represents each enriched functional term/pathway. Distance means the similarity between the connected functional terms. The figure was generated by ClueGO (*54*). C. ADAR2 enrichment on the *Malt1* transcript in Th17 cells as detected by RIP and RT-qPCR. * p-value<0.05 (t-test, n=3).

**Figure S4.**
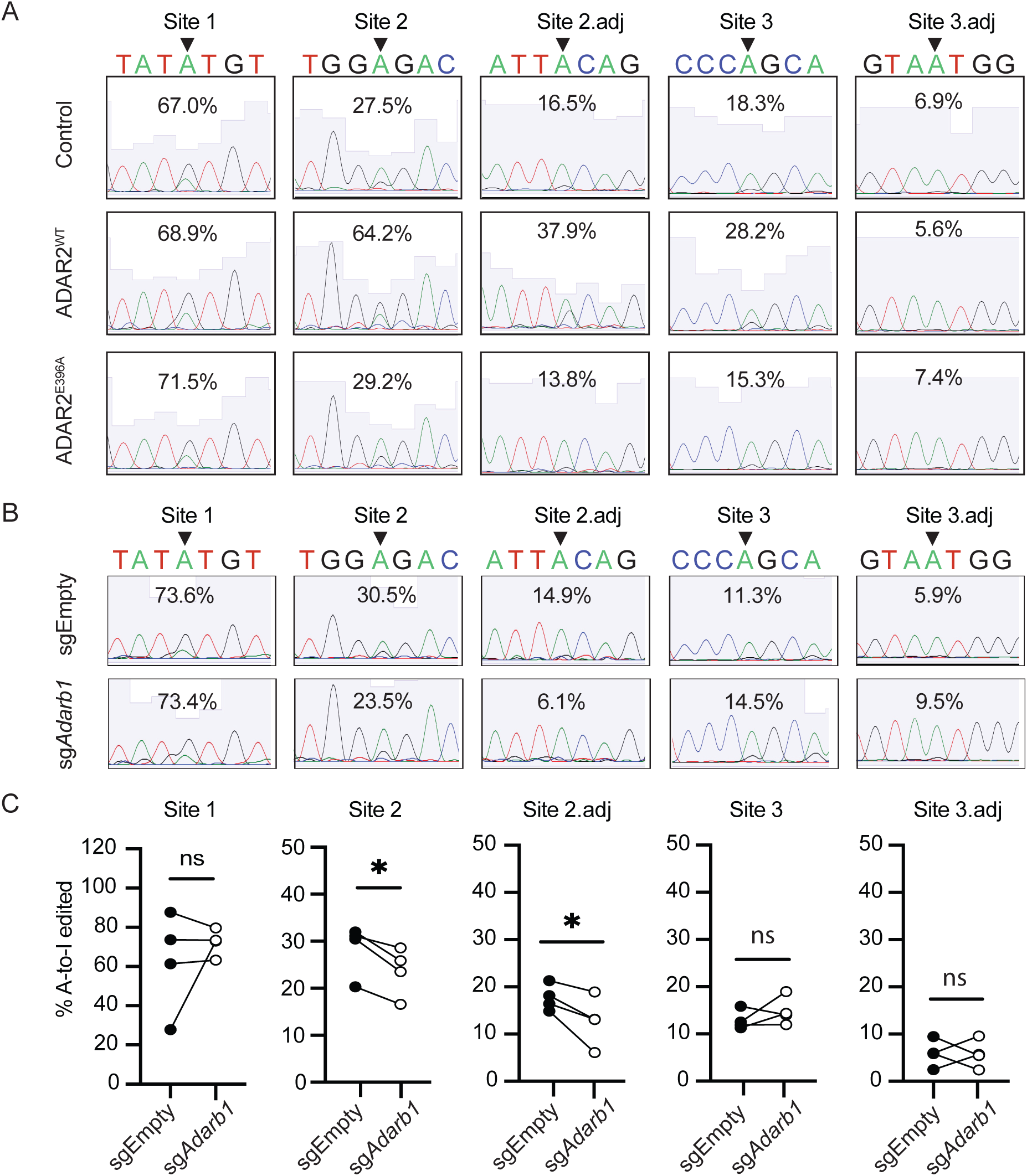
A-to-I edited sites on *Malt1* 3’UTR in culture Th17 cells. A. Representative sanger sequencing analyses of *Malt1* editing sites in Th17 cells transduced with the indicated expression vectors. B. Representative RT-PCR and Sanger sequencing analyses of *Malt1* editing sites in Cas9^+^ Th17 cells transduced with the indicated sgRNA expression vectors. C. Summarized editing proportion of the experiments described in B. Connecting lines indicate results from one independent experiment. * p-value<0.05, ns: not significant (paired t-test, n=4).

**Figure S5.**
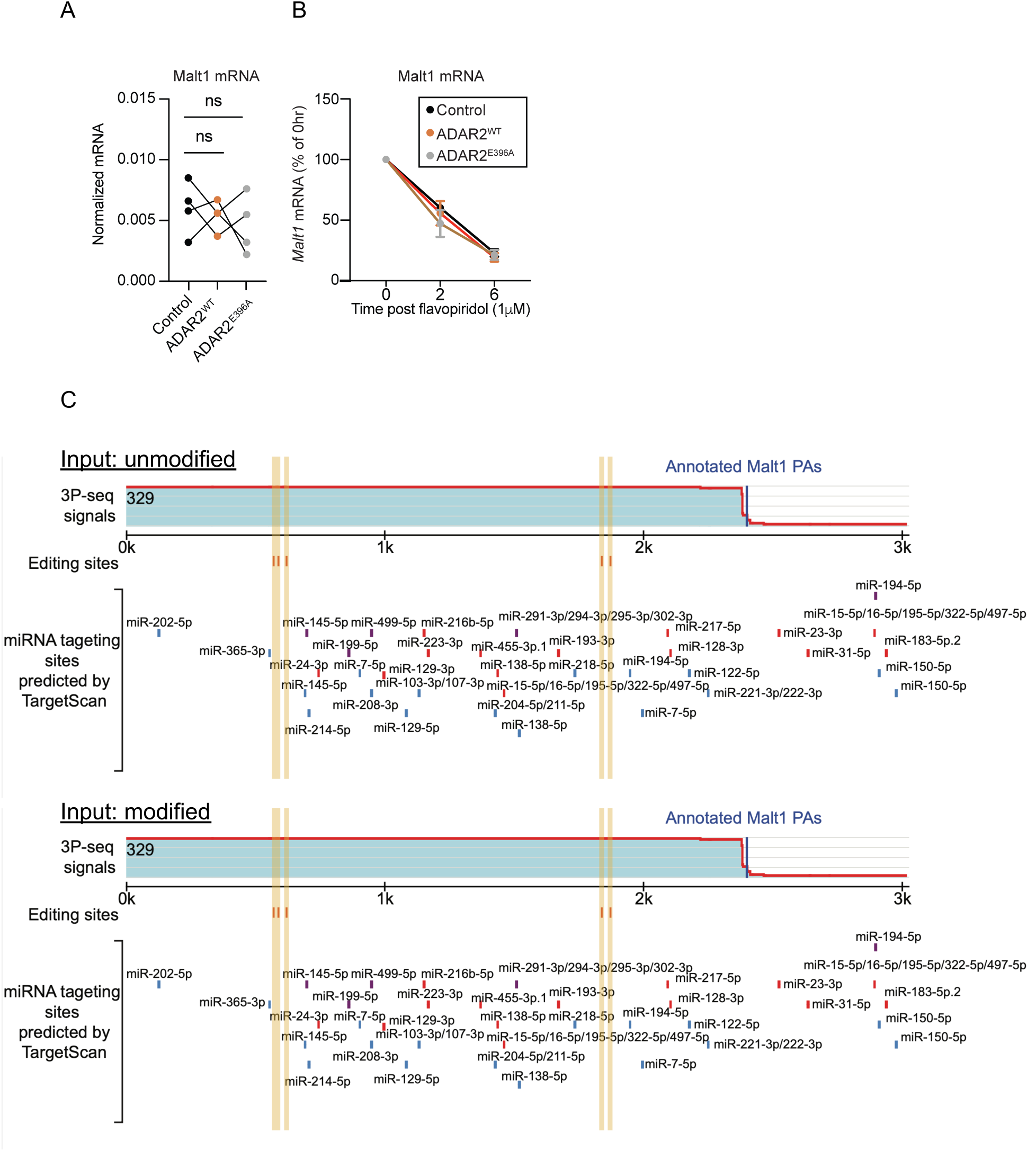
ADAR2 does not regulate *Malt1* transcription, RNA stability, and microRNA binding. A. Normalized expression of *Malt1* mRNA as assessed by qRT-PCR in Th17 cells transduced with the indicated vectors. *Gapdh* was used for normalization. Connecting lines indicate results from one independent experiment. ns: not significant (paired t-test, n=4). B. Relative *Malt1* mRNA level in Th17 cells transduced with indicated vectors treated with 1 μM flavopiridol for 0, 2, and 6 hrs. *Gapdh* was used for normalization. C. microRNA target sites on the unmodified (top) or A-to-I modified (bottom) *Malt1* 3’UTR predicted by TargetScan (*55, 56*).

**Figure S6.**
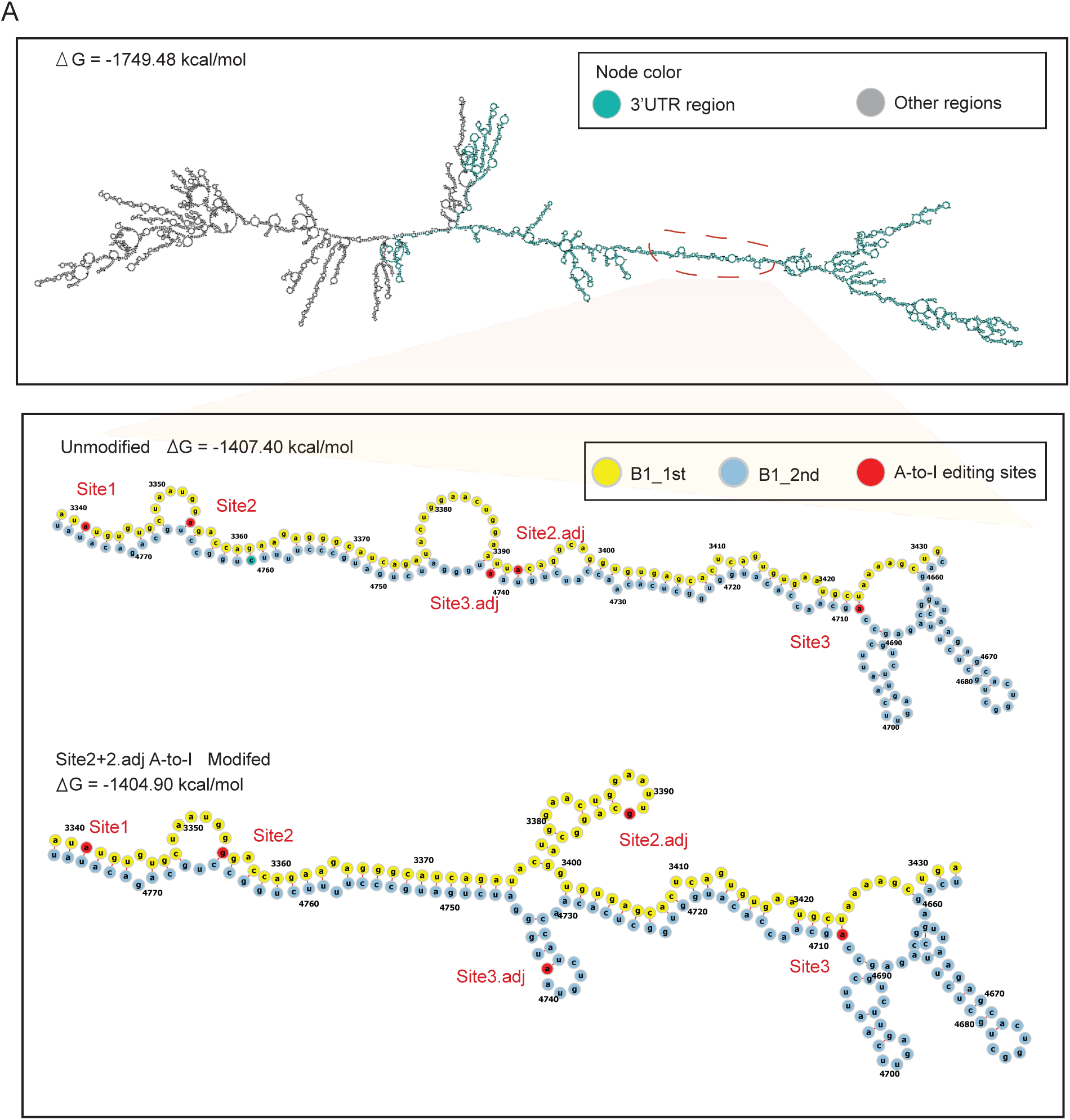
Predicted *Malt1* mRNA structures. A. Predicted RNA structures of the *Malt1* mRNA (top) and the 3’ UTR region harboring the two B2 SINE invert repeats that were unmodified (middle) or modified at sites 2 and 2.adj (bottom).

**Figure S7.**
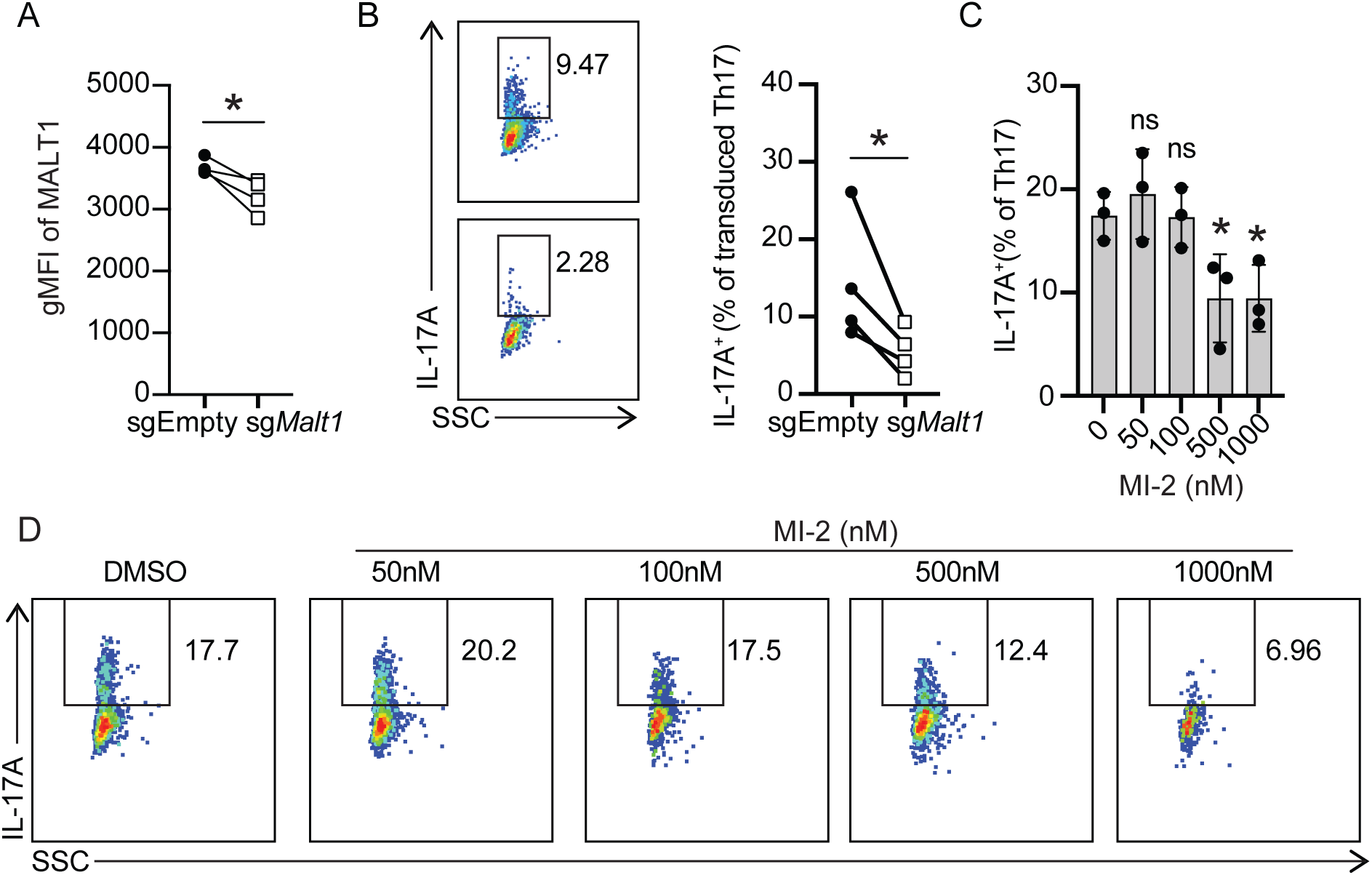
MALT1 induces Th17 cytokine production in a protease activity-dependent manner. A. Summarized gMFI of intracellular MALT1 in Cas9^+^ Th17 (CD90.1^+^RORγt^+^) cells transduced with the indicated vectors. Each line represents results from an independent experiment. * p-value<0.05 (paired t-test, n=4). B. Representative flow cytometry analysis (left) and summarized proportion of IL-17A^+^ Th17 cells from A (right). SSC: side scatter as an indicator of cell granularity. Each line represents results from an independent experiment. * p-value<0.05 (paired t-test, n=4). C. The proportion of IL-17A^+^ Th17 cells cultured in the presence or absence of the MALT1 inhibitor MI-2 at the indicated concentrations for 72 hrs. * p-value<0.05 (paired t-test with 0 nM, n=3). D. Representative flow cytometry analysis from C. SSC: side scatter as an indicator of cell granularity.

**Figure S8.**
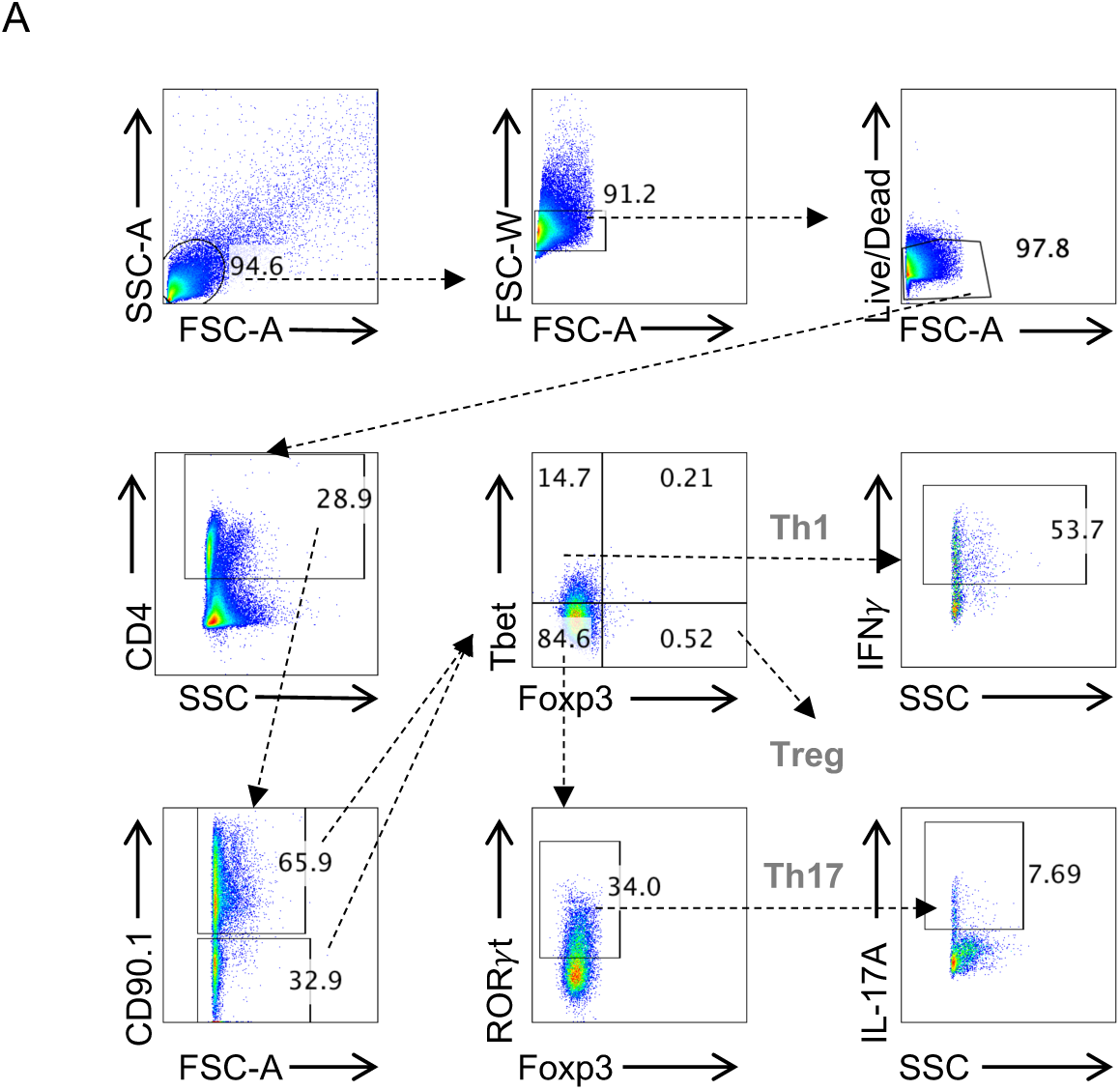
Gating strategy to examine colonic lamina propria T cells. A. Following the lymphocyte gate, doublets were eliminated, and live cells were analyzed for CD4 expression. Transduced CD90.1^+^CD4^+^ or non-transduced CD90.1^-^CD4^+^ T cells were divided into different T helper subsets according to Tbet, RORγt, and Foxp3 expression patterns. IFNγ and IL-17A expressions were further evaluated in each subset.

**Figure S9.**
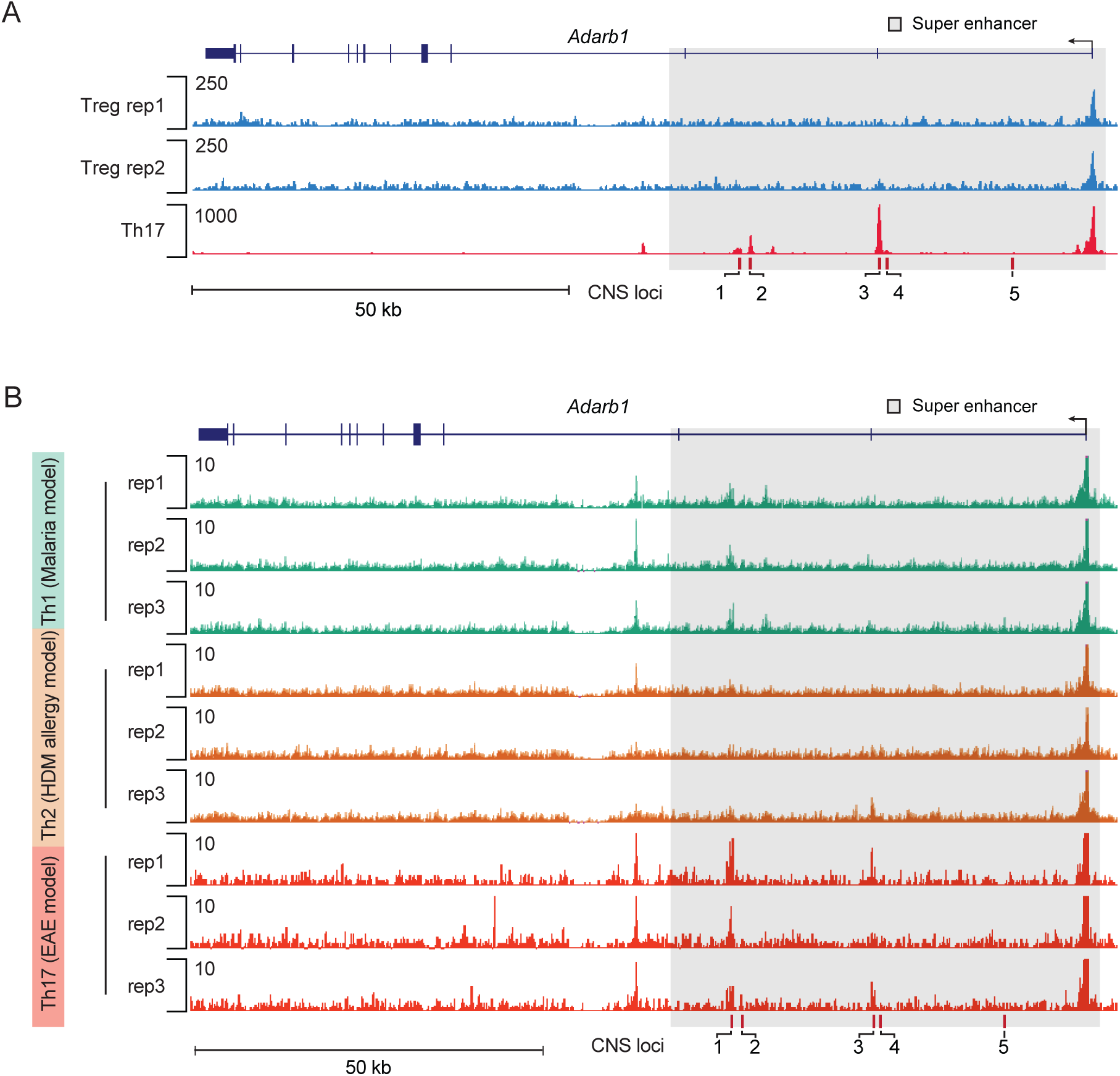
Chromatin accessibility at the *Adarb1* locus in different T helper subsets. A. Genome-browser tracks of ATAC-seq signals at the *Adarb1* locus in cultured Treg and Th17 cells (*42*). B. Genome-browser tracks of ATAC-seq signals at the *Adarb1* locus in Th1, Th2, and Th17 cells from malaria, house dust mite (HDM), or the experimental autoimmune encephalomyelitis (EAE) challenged mice (*41*).

**Figure S10.**
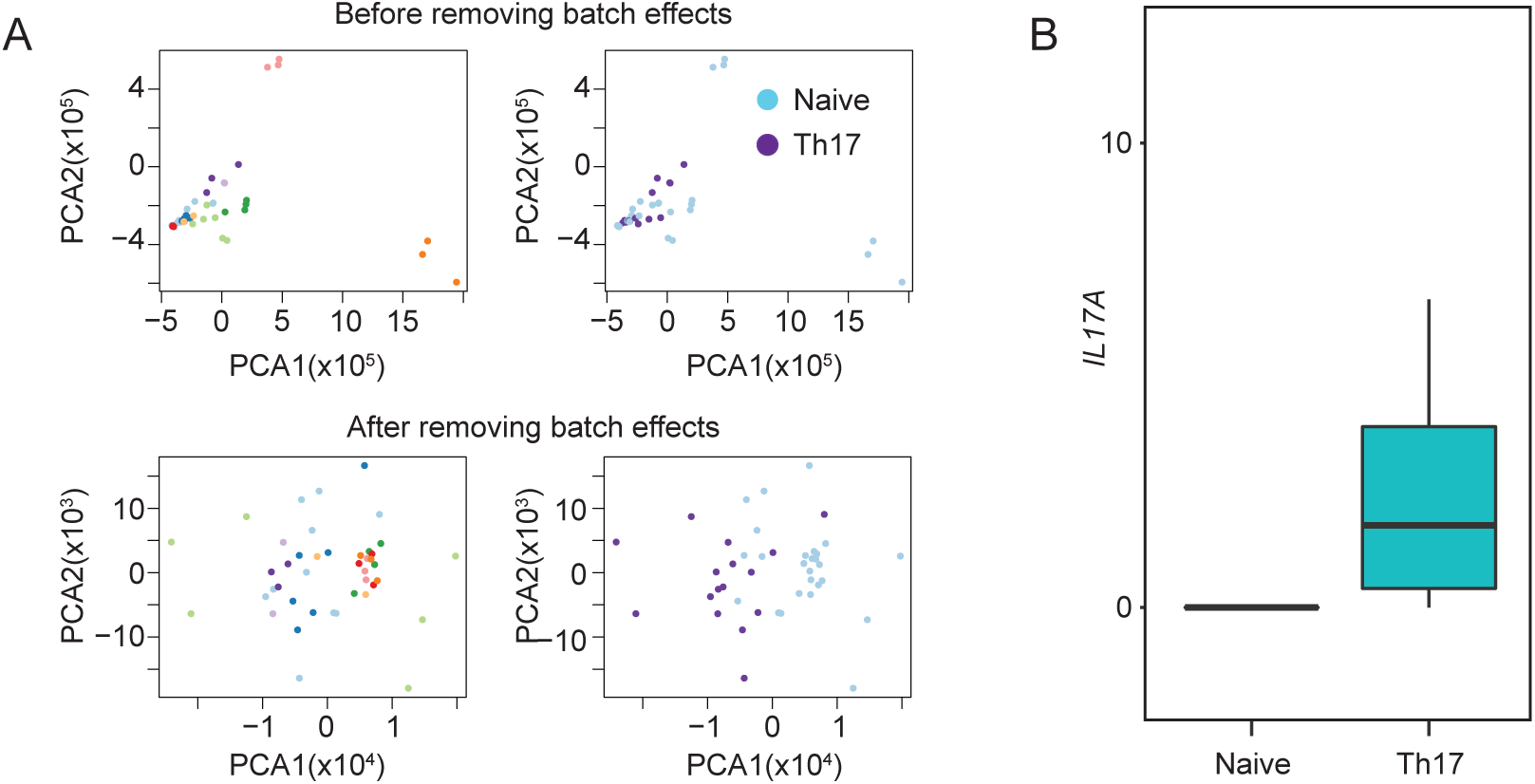
RNA-seq analysis of human naïve and Th17 cells. A. PCA analysis of 5 independent human RNA-seq datasets from naïve and culture Th17 cells before and after batch effect removal (*43*). B. Human *IL17A* expression level in naïve and culture Th17 cells as determined by RNA-seq from A (*43*).

**Figure S11.**
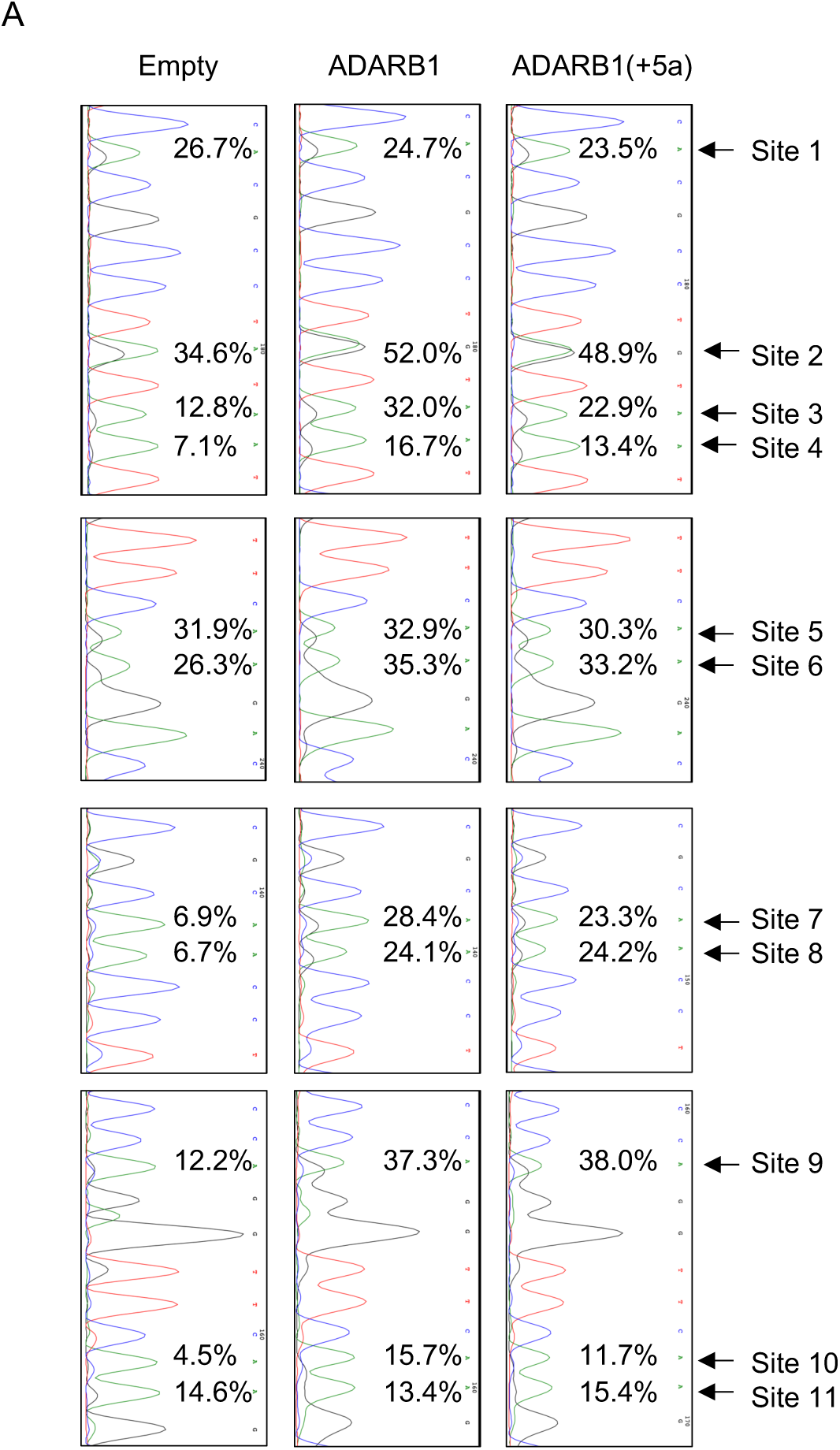
ADAR2 edits select sites on *MALT1* 3’UTR in HEK293 cells. A. Representative Sanger sequencing analyses of *Malt1* A-to-I modified sites in HEK293 cells transduced with the indicated vectors.

